# ZBTB33 (Kaiso) methylated binding sites are associated with primed heterochromatin

**DOI:** 10.1101/585653

**Authors:** Quy Xiao Xuan Lin, Khadija Rebbani, Sudhakar Jha, Touati Benoukraf

**Author notes:** Correspondence to: Touati Benoukraf, Ph.D. Faculty of Medicine, Discipline of Genetics, Craig L. Dobbin Genetics Research Centre, Room 5M317, Memorial University of Newfoundland St. John’s, NL A1B 3V6, Canada, Cancer Science Institute of Singapore National University of Singapore Centre for Translational Medicine, 14 Medical Drive, #12-01,Singapore 117599.

## Abstract

**Background:** ZBTB33, also known as Kaiso, is a member of the zinc finger and BTB/POZ family. In contrast to many transcription factors, ZBTB33 has the ability to bind both a sequence-specific consensus and methylated DNA. Although these dual binding preferences enable ZBTB33 to function as an active as well as repressive regulator of gene expression, little is known about the underlining molecular mechanisms.

**Results:** In this study, we aimed to investigate the role of ZBTB33 as a methylated DNA binding factor. We took advantage of the latest releases of the ENCODE sequencing datasets, including ZBTB33 ChIP-seq, whole genome bisulfite sequencing (WGBS), histone mark ChIP-seq and sequencing assays determining the chromatin states, to characterize the chromatin landscapes surrounding methylated ZBTB33 binding sites. Interestingly, our integrative analyses demonstrated that the majority of methylated ZBTB33 binding sites were located within condensed chromatin, which are inaccessible to DNase I and Tn5 transposase. Moreover, these sites were carrying a newly revealed histone post-translational modification signature, with significant enrichment of mono-methylation at lysine 4 of histone 3 (H3K4me1) and a complete absence of other active or expected repressive histone marks.

**Conclusions:** Overall, our analyses revealed that ZBTB33 has the unique ability to bind methylated DNA across heterochromatin in a transition state, suggesting a potential role for ZBTB33 in heterochromatin priming.

## Background

ZBTB33, also known as Kaiso, is a member of the zinc finger and BTB/POZ family. Similar to DNA (cytosine-5)-methyltransferase 1 (DNMT1) depletion, ZBTB33 knockdown phenotypes in *Xenopus* embryos include precocious activation of gene expression, apoptosis and developmental arrest [1]. Likewise, ZBTB33 depletion in the K562 cell line inhibits granulocyte differentiation and promotes cell proliferation [2]. In mammals, though, ZBTB33-knockout mice lack any significant phenotypes [3,4]; the depletion of ZBTB33 leads to intestinal tumorigenesis resistance [3], splenomegaly [4], increased locomotion and reduced volume of the lateral ventricles [5].

Like other BTB/POZ proteins, ZBTB33 forms a homodimer or heterodimer via its highly conserved BTB/POZ domains [6,7]. In contrast to many transcription factors, ZBTB33 has the ability to bind both a sequence-specific consensus and methylated DNA [8,9], which allows ZBTB33 to act as an activator or repressor of gene expression. In many types of cancer cells, ZBTB33 methyl-CpG-dependent binding events in the proximal promoters of essential tumor suppressor genes, such as Cyclin-dependent kinase inhibitor 2a (Cdk2a), Hypermethylated in cancer 1 (H1c1) and O-6-methylguanine-DNA methyltransferase (Mgmt) [10], are associated with repressive regulation, suggesting a crucial role in tumorigenesis. Further, ZBTB33 binding activity within the methylated promoter of E-cadherin is associated with E-cadherin silencing and an increase in cancer cell epithelial-mesenchymal transition (EMT) [11]. Additionally, ZBTB33 is capable of downregulating tumor suppressor microRNAs, such as miR-31 [12,13], in malignant prostate cancer cell lines. Interestingly, the repression of miR-31 appears to be methylation dependent, as it can be abrogated by demethylation of the ZBTB33 binding site *via* 5-aza-2’-deoxycytidine (5-aza-dC) treatment [12].

The advent of whole genome sequencing combined with DNA bisulfite treatment brought to light a myriad of consequences of DNA methylation in gene regulation and chromatin modulation. Indeed, methylation of cytosine is not solely associated with gene repression but also with RNA splicing, nucleosome positioning and transcription factor recruitment [14]. These events might be mediated by proteins such as ZBTB33, which can recognize and bind a specific methylated DNA motif. In this study, we aim to investigate the role of ZBTB33 as a methylated DNA binding factor. We took advantage of the latest release of the ENCODE sequencing datasets, including ZBTB33 ChIP-seq, whole genome bisulfite sequencing (WGBS), histone mark ChIP-seq and sequencing assays determining the chromatin states, to characterize the chromatin landscape in methylated ZBTB33 binding sites.

## Results

### ZBTB33 binding sites are predominantly methylated

Although ZBTB33 has been shown to bind numerous methylated loci, the genome-wide generalization of this ability remains unclear. By comparing a previous release of ENCODE ZBTB33 ChIP-seq dataset with WGBS and reduced representation bisulfite sequencing (RRBS) in GM12878 and K562 cell lines, respectively, a recent study by Blattler *et* al. concluded that ZBTB33 binds predominantly non-methylated DNA [15]. However, the ChIP-seq datasets used in these analyses were sequenced at a low depth of coverage in both cell lines (36nt single-end read and less than 20 million reads in each replicate, Supplementary Table 1 and 2) and, the antibody quality has not fully fulfilled the ENCODE Consortium standards while absolved by an ENCODE Antibody Review Committee (Supplementary Table 1 and 2, ENCODE Antibody ID: ENCAB000AML). These ChIP-seq experiments revealed a low number of ZBTB33 binding sites in both cell lines (2,496 peaks in GM12878 cells and 2,740 peaks in K562 cells) and poor consistency in replicate assays performed in K562 cells (Supplementary Table 1 and 2). Moreover, due to the limitation of genome sequencing coverage in RRBS experiments, the characterization of the DNA methylation status of ZBTB33 binding sites in K562 cells was restricted to approximately 4% of CpG sites in the genome, primarily located in gene promoters and CpG islands [16]. To extend Blattler *et* al. study, we performed methylation profiling of ZBTB33 binding sites in K562 using WGBS, which also revealed hypomethylated ZBTB33 bound promoters, while a small portion of hypermethylated ZBTB33 binding sites located in introns (Supplementary Figure 1).

The latest ZBTB33 ChIP-seq datasets in K562 and GM12878 cells released by ENCODE have shown a significant increase in antibody quality (both antibody primary and secondary tests are compliant to the current ENCODE standards, ENCODE Antibody ID: ENCAB292USO), library quality and reproducibility (Supplementary Tables 2-4, Supplementary Figure 2), encouraging us to perform a novel ZBTB33 binding sites / DNA methylation integrative analysis to determine the methylation status of ZBTB33 binding sites. In this K562 ChIP-seq release, we identified approximately nine times more ZBTB33 binding sites (25,170 peaks) compared to the previous experiment (2,740 peaks), with a slight overlap between the two experiments (Supplementary Figure 3). Finally, the overall methylation status for ZBTB33 narrow peak regions were profiled as hypermethylated in K562, while in GM12878 the CpG sites within ZBTB33 peaks can be either hyper- or hypomethylated (Supplementary Figure 4). Our analysis shows a different DNA methylation profile compared to the Blatter *et* al. results, which is mainly due to the distinct ZBTB33 peak sets identified between the two studies (Supplementary Figure 3). However, given the improved quality in the latest release of the ZBTB33 ChIP-seq in both cell lines (Supplementary Table 2), we suggest that peak sets derived from our analysis better describe the genome-wide occupancy of ZBTB33 binding sites. That is the reason why we decided to perform a new integrative analysis of ZBTB33 binding sites with epigenomic features using the latest ENCODE datasets in both cell lines.

We performed a *de novo* motif discovery using MEME-ChIP [17] and *peak-motifs* module from RSAT package [18] in K562 and GM12878 (Supplementary Figure 5). in K562 and GM12878 (Supplementary Figure 5). Similar motifs were discovered in K562 and GM12878 using the same tool, while the motif discovery results in both cell lines were not consistent between MEME and RSAT. The difficulty in *de novo* motif discovery may be due to the atypical intrinsic sequences surrounding ZBTB33 peak centers, overrepresented in CG and GC dinucleotides while underrepresented in other dinucleotide contents in both cell lines (Supplementary Figure 6). Therefore, we decided to rely on the ZBTB33 motif TCTCGCGAGA described and validated by Raghav *et* al. [19]. Indeed, in this study, the authors proved high binding affinities of ZBTB33 to the methylated palindromic sites *in vitro*, even higher than the sequence-specific consensus Kaiso binding site (KBS) TCCTGCNA, and further showed in detail that methylation in the core CGCG is crucial for the binding affinity while the flanking sequences play an important role. Consequently, we searched for the known ZBTB33 binding motif (TCTCGCGAGA) [19] with a position weight matrix downloaded from JASPAR (JASPAR ID: MA0527.1) across all identified peaks using the *matrix-scan* module from RSAT [20]. The ZBTB33 binding motif TCTCGCGAGA was found to be centrally enriched around ZBTB33 peak centers in both cell lines, while the top three motifs discovered by MEME were either not as centrally enriched as TCTCGCGAGA or not conserved across cell lines (Supplementary Figure 7). Further spatial correlation analyses between the genome-wide TCTCGCGAGA sites and ZBTB33 peaks by GenometriCorr [21] and regioneR [22] revealed a significantly high overlap between the two genomic intervals in both cell lines, corroborating a high enrichment of the known ZBTB33 motif in the peak regions (Supplementary Table 5-6 and Supplementary Figure 8). At last, more than 60% of the identified peaks contained the ZBTB33 binding motif in both cell lines (Figure 1A, Supplementary Table 7), confirming the validity of the ChIP-seq experiments. Overall, ZBTB33 binding sites were located mostly within introns or intergenic regions in terms of peak number (Supplementary Figure 9). However, when normalized according to the length of different genomic features, ZBTB33 binding sites were revealed to be more enriched in gene promoters in both cell lines, suggesting a direct role in gene regulation (Supplementary Table 8 and 9). DNA methylation levels within ZBTB33 binding sites were not distributed homogenously across the genome. Indeed, binding sites in intronic and intergenic regions were found to be highly methylated, while in promoter loci, a considerable portion of CpG sites bound by ZBTB33 were unmethylated (Figure 1B). As described previously [9], our analysis confirms that ZBTB33 has the ability to bind both methylated and unmethylated DNA. When comparing ZBTB33 binding properties in the two cell lines, we observed that most of ZBTB33 binding sites were not conserved across cell lines (16,272 out of 16,954 and 1,375 out of 2,057 ZBTB33 binding sites are cell specific in K562 and GM12878 respectively - Supplementary Figure 10). Interestingly, the 682 ZBTB33 binding sites common to both cell lines shared similar DNA methylation patterns, with a high level of methylated ZBTB33 binding sites, particularly located within intronic peaks (Supplementary Figure 10). The main difference that we can observe across these two cell lines is a distinct methylation profile on ZBTB33 bound promoters, which are more hypermethylated in K562 and hypomethylated in GM12878. This difference is probably due to a distinct chromatin and epigenetic landscape across cell lines (Chronic myeloid leukemia cell - K562 versus B lymphocyte - GM12878), which leads to different binding properties.

**Figure 1.** Characterization of ZBTB33 binding sites. **A)** Venn Diagram represents our ZBTB33 peak selection in K562 and GM12878. Among all IDR peaks discovered (blue area), motif scanning identified peaks containing a ZBTB33 motif (orange area). Further, we captured the ZBTB33 binding sites which were covered by the ENCODE WGBS, by at least 5 reads in each of two replicates in K562 (i.e., 5 reads X 2, gray area) and 5 reads in one replicate in GM12878 (i.e., 5 reads X 1, gray area). Then, we characterized 3,522 binding sites as methylated (purple area) while 266 were unmethylated (yellow area) in K562, and 802 binding sites as methylated (purple area) while 505 were unmethylated (yellow area) in GM12878. **B)** Distributions of CpG beta scores within ZBTB33 binding sites according to different genomic location contexts. **C)** Occurrence of the ZBTB33 binding motif within the ±2 kb regions surrounding ZBTB33 methylated and unmethylated ChIP-seq peaks computed by *matrix scan* from RSAT. In K562, to rule out number bias, (3,522 *vs* 266 binding sites), we randomly selected 266 binding sites across methylated ZBTB33 peaks for our comparison. The significance of the motif enrichment is determined by comparing the ZBTB33 motif enrichment with 3 permutated motifs derived from the ZBTB33 binding motif matrix. **D)** Logo represents the ZBTB33 position-specific weight matrix along with beta scores on CpG sites. Unmethylated CpG (beta score less than 10%) are represented in blue, methylated (beta score greater than 90%) in orange and heterogeneously methylated sites in green. Numbers on the top of bars represent the amount of sequences containing a CpG covered by WGBS.

To further investigate the chromatin landscapes of methylated and unmethylated ZBTB33 binding sites, we considered all CpG sites within the motif that had a methylation level greater than 90% as methylated binding sites. Conversely, unmethylated binding sites were defined as the loci where CpG methylation levels were less than 10%. These stringent thresholds allowed us to avoid mis-scoring DNA methylation on ZBTB33 binding sites due to cell heterogeneity. In other words, choosing extreme cut-offs helped us to focus our analysis only on loci highly homogenous in term of DNA methylation. With these thresholds and settings described in the methods section, we identified 3,522 highly methylated ZBTB33 binding sites and 266 unmethylated sites in K562, and 802 highly methylated ZBTB33 binding sites and 505 unmethylated sites in GM12878 (Figure 1A). While the ZBTB33 binding motif was significantly enriched in both methylated and unmethylated binding sites (p-value < 2.2 10^-16^, Fisher’s exact test), we observed a significant central enrichment of ZBTB33 motif in methylated sites compared to unmethylated sites (p-value < 10^-5^, Fisher’s exact test, Figure 1C). This observation suggests that ZBTB33 ChIP-seq peaks found within methylated DNA loci contain more ZBTB33 direct binding sites compared to peaks detected across unmethylated DNA loci [23]. Interestingly, the ZBTB33 binding motif showed a high level of conservation in these methylated CpG sites, implying a strong selection pressure for these bases (Figure 1D). Finally, we found that the sequence-specific consensus KBS TCCTGCNA, previously characterized by *in vitro* assays [8], was not enriched neither in ZBTB33 K562 nor GM12878 ChIP-seq peaks (Supplementary Figure 11).

### ZBTB33 binding domains are associated with genes involved in chromatin regulation

We annotated both methylated and unmethylated ZBTB33 binding sites according to their respective chromatin contexts and functions using the previous systemic annotations performed in K562 and GM12878 by the Broad Institute (ChromHMM: Chromatin state discovery and characterization) [24]. Most methylated ZBTB33 binding sites were associated with weak transcription functions, including weak promoter, weak/poised enhancer and weak transcription (Supplementary Figure 12 left). However, annotation results for unmethylated ZBTB33 binding sites were correlated with more transcriptionally active regions, with higher proportions annotated as active promoters and strong enhancers (Supplementary Figure 12 right). Notably, a considerable portion of unmethylated ZBTB33 binding sites were located within insulator regions (Supplementary Figure 12 right), corroborating that ZBTB33 is an interacting partner of CTCF [25]. In addition, gene ontology (GO) analysis of target genes of ZBTB33 methylated regulatory elements computed by the Genomic Regions Enrichment of Annotation Tool (GREAT) [26], revealed a significant enrichment of terms relative to epigenetic regulation of gene expression, nucleosome organization and histone lysine methylation in K562 (Figure 2), which suggests an important role of ZBTB33 in chromatin modulation. The limited number of peaks in ZBTB33 ChIP-seq experiment in GM12878 did not allow us to confirm this results in this cell line. In contrast, no significant GO term was found for unmethylated binding sites, probably because of the low number of loci.

**Figure 2.** Gene ontology of genes targeted by methylated ZBTB33 binding sites in K562. This bar plot represents the -log10 transformation of FDR values for each GO term enriched in ZBTB33 targeting genes in K562 calculated by GREAT software. There were no results for unmethylated ZBTB33 binding sites.

### Co-occupancy of transcription factors surrounding ZBTB33 binding sites

To identify the potential protein complex associated with ZBTB33, we analyzed K562 and GM12878 ENCODE ChIP-seq profiles, including 187 and 114 transcription factors with more than 30 million uniquely map reads in K562 and GM12878 respectively, surrounding methylated and unmethylated ZBTB33 binding sites. For all these transcription factors, only one significant binding event (MLLT1) was detected surrounding methylated ZBTB33 binding sites in both cell lines (Figure 3 left). Interestingly, MLLT1, also known as ENL, is a critical component of the super elongation complex and also a histone acetylation reader [27]. The co-occupancy of MLLT1 with ZBTB33 in methylated binding sites suggests the potential of transcriptional regulation in these regions. No significant signal was found surrounding methylated ZBTB33 binding sites within the ENCODE ChIP-seq datasets of transcription factors sequenced with less than 30 million uniquely map reads (34 for K562 and 18 for GM12878, Supplementary Figure 13 left). This analysis shows that, within methylated DNA, ZBTB33 does not commonly exit as a member of a protein or regulatory complex but presumably binds to its DNA targets solitarily in most instances, a feature that ZBTB33 shares with pioneer transcription factors [28]. In contrast, considerable co-occupying events were detected surrounding unmethylated ZBTB33 binding sites (Figure 3 right, Supplementary Figure 13 right). Hierarchical clustering on these TF ChIP-seq signals surrounding unmethylated ZBTB33 binding sites clearly identified three distinct clusters: (1) TFs that co-bind with ZBTB33, (2) TFs that do not co-bind with ZBTB33 and (3) TFs that co-bind with ZBTB33 with a higher affinity (Figure 3 right). Remarkably, in K562, the cluster 3 included the histone lysine demethylase PHF8, suggesting a role for ZBTB33 in removing the di-methyl group of lysine 9 in histone H3 (H3K9me2) [29,30]. In addition, the fact that MAX and RBFOX2 belonged to cluster 3 in K562 indicates that ZBTB33 may be included in protein complexes regulating cell proliferation as well as apoptosis and EMT [31,32]. Unfortunately, PHF8 and RBFOX2 ChIP-seq datasets were not available for GM12878. However, ZBTB33-MAX co-binding was confirmed in GM12878. Finally, ZBTB33 binding sites also co-localized with the insulator protein CTCF (cluster 1 for K562 and cluster 3 for GM12878), and further analyses using the TFBS cluster database from UCSC (*c.f.* methods section) confirmed that approximately 29% and 43% of ZBTB33 binding sites co-localized with CTCF in K526 and GM12878 cells respectively, corroborating a previous study [25] (Supplementary Figure 14, Supplementary Tables 10 and 11).

**Figure 3.** Co-occupancy of 187 and 114 transcription factors surrounding un/methylated ZBTB33 binding sites in K562 and GM12878 respectively. Heatmaps describe the enrichment (average normalized read count) of 187 and 114 transcription factors from ENCODE datasets (ChIP-seq containing more than 30 million uniquely mapped reads) within ±500 bp regions surrounding methylated and unmethylated ZBTB33 binding sites in K562 **(A)** and GM12878 **(B)** respectively. Here, each row represents a sequencing experiment from ENCODE where read intensities for all ZBTB33 peak regions were averaged, while each column represents relative distance to the ZBTB33 peak center. Additional heatmaps portraying CTCF, RBFOX2, PHF8 and MAX ChIP-seq signals surrounding unmethylated ZBTB33 binding sites in K562 (A, right side) and MAX and CTCF signals surrounding unmethylated ZBTB33 binding sites in GM12878 (B, right side) are highlighted. In these highlighted heatmaps, each row represents a ZBTB33 binding site region, and each column represents the relative distance from the ZBTB33 ChIP-seq peak center. Furthermore, the fold enrichment of the average normalized read counts in the peak regions (±100 bp around peak centers) compared to the flanking regions (200 bp windows located 300 bp away from the peak centers) for each TF is shown in a heatmap. Orange color represents the average normalized read count in the peak regions is more than in the flanking regions, while the blue color denotes the opposite.

### ZBTB33 binds predominantly heterochromatin loci harboring a primed chromatin histone signature

To characterize chromatin landscapes associated with ZBTB33 binding events in conjunction with DNA methylation levels, we integrated ZBTB33 ChIP-seq results with ENCODE datasets from various histone modifications, RNA polymerase II activities and chromatin states in K562 and GM12878 cells. Unmethylated ZBTB33 binding sites harbored an expected histone code, with enrichment of active chromatin marks (H3K4me1, H3K4me3 and H3K27ac), depletion of repressive histone marks (H3K9me3 and H3K27me3), active polymerase II transcription, and open chromatin footprints revealed by formaldehyde-assisted isolation of regulatory elements (FAIRE-seq), DNase l hypersensitive site sequencing (DNase-seq) and assay for transposase-accessible chromatin (ATAC-seq) experiments (Figure 4, Supplementary Figure 15 and 16). However, the total absence of FAIRE-seq, DNase-seq and ATAC-seq signals across methylated ZBTB33 binding sites suggests that these domains are located within the heterochromatin. Intriguingly, these heterochromatin domains bound by ZBTB33 carried a unique post-translational histone modification combination, with a high enrichment of H3K4me1 and total absence of other active and repressive histone marks (Figure 4, Supplementary Figures 15 and 16). This observation is consistent in both cell lines. In order to investigate whether such a unique chromatin landscape is associated with ZBTB33 or just correlated with DNA methylation, we profiled the signal intensities of H3K4me1 ChIP-seq in a 5bp window size around all CpGs found in methylated ZBTB33 binding sites (cf. Methods). Furthermore, as controls, we used the same strategy to profile CpGs in unmethylated ZBTB33 binding sites and all sequenced methylated CpGs in the genome (Supplementary Figure 17). Our analysis clearly shows three states of H3K4me1 enrichment in K562: 1) low, which is associated with methylated CpGs only; 2) high, which is associated with unmethylated CpGs in ZBTB33 binding sites and 3) intermediate, which is associated with methylated CpGs found in ZBTB33 binding sites. In GM12878, H3K4me1 enrichment surrounding methylated CpGs found in ZBTB33 binding sites is higher compared to unmethylated ZBTB33 associated CpGs, which is due to a much higher portion of unmethylated ZBTB33 binding sites located in promoters compared to the methylated sites (Supplementary Figure 9B and C). Therefore, this analysis provides an additional evidence on the association between ZBTB33 and primed heterochromatin. H3K4me1 enrichment is known to mark poised enhancers, as well as active enhancers when it is coupled with H3K4me3 and/or H3K27ac [33,34]. Nevertheless, thus far, H3K4me1 has never been described as being associated with heterochromatin. Here, methylated ZBTB33 binding loci introduce a novel chromatin state combining compacted DNA, accessibility to a transcription factor, and histone post-translational modifications in an intermediate state, between repressive and active chromatin (Figure 5).

**Figure 4.** Chromatin landscapes in un/methylated ZBTB33 binding sites. Heatmaps describe the DNA methylation profile (average beta score) as well as enrichment (average normalized read count) of histone marks, RNA polymerase II and sequencing datasets indicating chromatin states within the ±5 kb regions surrounding unmethylated (left side) and methylated (right side) ZBTB33 binding sites in K562 **(A)** and GM12878 **(B)**. Here, each row represents a sequencing experiment from ENCODE where CpG beta scores or read intensities for all ZBTB33 peak regions were averaged, while each column represents relative distance to the ZBTB33 peak center. Examples of chromatin landscapes surrounding unmethylated and methylated ZBTB33 binding sites in K562 are shown in panels **(C)**.

**Figure 5.** Proposed model of ZBTB33 methylated DNA-dependent binding events. The canonical heterochromatin state (top) is featured as a compact DNA structure deposited with H3K9me3 and H3K27me3, while an euchromatin state (middle) is characterized by an uncondensed nucleosome structure where H3K4me1/3 and H3K27ac marks are enriched. The fact that ZBTB33 is able to recognize and directly bind methylated and condensed DNA in a sequence-specific manner on DNA domains associated with H3K4me1 enrichment and depletion of repressive marks suggests that ZBTB33 may play a role in heterochromatin priming (bottom).

## Discussion

During the recent decades, DNA methylation has commonly been associated with transcriptional repression. However, this correlation is not as strict as previously assumed, and emerging iconoclastic evidence has revealed an increasing number of transcription factors with the ability to bind methylated DNA [35–37]. These observations suggest a more complex interplay between DNA gene expression and DNA methyl-modification that is dependent on the methylcytosine context. Accumulating evidence supports the hypothesis that TFs serve as methylated-CpG readers at enhancers or within regions of low CpG density, while methyl-CpG-binding domain proteins are prone to recognize densely methylated DNA sites [38]. ZBTB33 is one of the TFs which have been recognized with the ability to bind methylated DNA. It has been reported that at least two symmetrically methylated CpG dinucleotides are required to be bound by ZBTB33 *in vitro* [39], while a recent *in vitro* study showed that one methylated CpG is even necessary and sufficient for ZBTB33 recognition [40]. Indeed, the newly deposited ZBTB33 ChIP-seq datasets have revealed approximately 60% methylated ZBTB33 binding sites possess the methylated CGCG core motif in both cell lines, while the rest contain either only one methylated CpG site or more than one, but not consecutive CpGs. ZBTB33 that binds methylated DNA is primarily described as a transcription repressor [41]. Nevertheless, the mechanism of the repression exerted by ZBTB33 when binding methylated DNA is still not well depicted. Yoon *et* al. proposed a model in which ZBTB33 is associated with nuclear receptor corepressor (N-coR), to form a “Kaiso/N-coR” complex and bind methylated promoters to foster histone deacetylation and, consequently, the formation of repressive chromatin structures [42]. However, at the genome scale, our analyses in K562 and GM12878 cells do not confirm the co-occupancy of N-coR and ZBTB33 across methylated DNA. This mechanism might be cell cycle specific and hardly detectable *via* ChIP-seq assays performed in an unsynchronized cell population. Interestingly, ZBTB33 is also able to function as a methyl-dependent activator [43]. A more recent work by Zhenilo *et* al. suggests that SUMOylation in ZBTB33 BTB/POZ protein-protein interaction domain may switch ZBTB33 from a gene repressor to an activator [44].

Overall, our integration of 221-TF and 132-TF ChIP-seq datasets in K562 and GM12878 cells respectively, with their corresponding methylome profiles suggests the two distinct functions of ZBTB33 are dependent of the DNA methylation context. When bound to unmethylated DNA, ZBTB33 co-localizes with co-factors crucial for the chromatin topology and histone tail modifiers, such as CTCF and PHF8, respectively. On the other hand, the majority of methylated ZBTB33 binding sites are located within compacted and methylated DNA. Therefore, we propose that ZBTB33 has the ability to recognize and bind a specific motif within methylated and compacted DNA localized in heterochromatin. More interestingly, these binding sites are associated with an enrichment of H3K4me1 alone, the absence of other active signatures (H3K4me3 and H3K27ac) and a depletion of the expected repressive marks (H3K9me3 and H3K27me3), suggesting that the targeted regions are poised enhancers [45,46]. Indeed, H3K4me1 was termed “a window of opportunity” by Eliezer and Joanna [47] in which enhancer activation is possible, followed by nucleosome removal and H3K27ac deposition. This function is reminiscent of territory occupied by pioneer factors, which uncompact the heterochromatin and initiate the recruitment of the transcriptional machinery [28]. However, further investigations are required to fully categorize ZBTB33 as a pioneer factor. Nonetheless, our analyses have illuminated new evidence supporting ZBTB33 priming of heterochromatin and potentially playing a role in the modulation of chromatin topology.

## Conclusions

Altogether, our bioinformatics analysis has discovered a unique role of ZBTB33 and this is through its ability to bind a specific motif within methylated and compacted DNA. Remarkably, ZBTB33 methylated binding sites are associated with a distinctive histone signature, where only H3K4me1 appears to be enriched. Although ZBTB33 binding sites are clearly located within condensed nucleosome loci, the histone code associated with ZBTB33 binding sites is incompatible with a canonical heterochromatin state, which requires a H3K9me3 enrichment to associate the HP1-Lamin B1 receptor complex [48]. Therefore, we suggest that ZBTB33 is involved in the inducement or the maintenance of a new characterized chromatin state, transitional between repressed and active form. Moreover, we found that H3K4me1 marks primed DNA loci not only in euchromatin as previously described [49], but also in heterochromatin.

## Methods

### ZBTB33 ChIP-seq peak calling

ZBTB33 ChIP-seq dataset as well as K562 and GM12878 DNA input files were downloaded from the ENCODE consortium website (https://www.encodeproject.org, Supplementary Table 1 and 3) and reads in raw files were aligned to the hg38 genome assembly using STAR with the following parameters: -- outFilterMultimapNmax 1, --outFilterMatchNminOverLread 0.80 and blocked splice function [50]. For peak calling, we used MACS2 [51] with default parameters and the irreproducible discovery rate (IDR) pipeline, as recommended by the ENCODE consortium [52], to assess the consistency between replicates.

### Motif identification surrounding peak regions

Motif *de novo* discovery was performed using MEME-ChIP package with default parameters [17] in a ± 100bp window size surrounding peak summits and *peak-motifs* module in RSAT package with oligo, position and local-word analyses [18] in a ± 500bp window size surrounding peak summits. Further, the RSAT motif discovery results underwent motif clustering using *matrix-clustering* module with default parameters [53].

For the known ZBTB33 motif scan, extended peak sequences (±4 kb surrounding peak summits) were scanned with the ZBTB33 position-specific weight matrix (PSWM) downloaded from JASPAR [54] (JASPAR ID: MA0527.1) using the RSAT *matrix-scan* module with a Markov model of order 1 as a background model, and a p-value threshold set to 10^-3^ [20,55]. Our illustrations highlight the significance of the motif enrichment by comparing the ZBTB33 PSWM with the randomly permuted motif matrices derived from the ZBTB33 PSWM. Direct ZBTB33 binding sites were identified by segregating ChIP-seq peaks with a ZBTB33 PSWM match within ±100 bp surrounding peak summits (Supplementary Table 7). For peaks with multiple motif matches, we selected the motif having the highest weight score and the nearest distance to peak summits. Similar strategy was applied to the other motif scans in the study.

Spatial correlation between genome wide potential ZBTB33 binding sites, identified using aforementioned motif scan protocol, and ZBTB33 peaks was confirmed using GenometriCorr with 100 permutation tests [21] and regioneR with 1000 overlap permutation tests [22].

### Methylation states in ZBTB33 binding sites

To observe DNA methylation profiles at ZBTB33 binding sites, whole genome bisulfite sequencing (WGBS) raw datasets in K562 and GM12878 were downloaded from the ENCODE consortium (Supplementary Table 12). Using Trim Galore and cutadapt [56], reads were trimmed according to fastQC results, and adaptors were also removed. Alignment and methylation extraction were achieved with bismark [57] using bowtie2 [58] and default parameters. In K562, after confirming a high correlation coefficient across replicates using methylKit [59] (Supplementary Table 12), we merged the replicated WGBS data to increase the sequencing coverage. Methylation levels of CpGs with at least at least 5x coverage in each replicate (i.e., 5 reads X 2) [60] located within ZBTB33 binding motifs were extracted with custom python scripts. DNA methylation (beta score) of ZBTB33 binding sites was calculated by using the ratio between all methylated-cytosine read number to total read number within the region, across both replicates as described previously by Adusumalli *et* al. [61]. We considered a region with a beta score greater than 90% as a homogeneously methylated binding site and a region with a beta score less than 10% as a homogeneously unmethylated binding site, while a binding site with an in-between beta score (more than 10% and less than 90%) is considered heterogeneously methylated across the cell population. In GM12878, we were constrained to use only one replicate due to the low amount of ZBTB33 ChIP-seq peaks. Indeed, only 789 peaks are covered by 5 reads X 2. Among them, 317 are methylated (beta score greater than 90%) and 160 unmethylated (beta score less than 10%). Methylation levels of CpGs within ZBTB33 binding sites in GM12878 cells were computed as explained previously on loci covered by a least 5 reads in one replicate (i.e., 5 reads X 1, Supplementary Table 12). We used the Fisher’s exact test to support that the occurrences of ZBTB33 motif matches surrounding methylated and unmethylated ZBTB33 peaks (100 bp around peak centers) were significantly higher than the occurrences of permuted motifs (control). Furthermore, to validate a more central enrichment of ZBTB33 motif surrounding the methylated sites compared to the unmethylated loci, we assessed the difference in proportions, between the methylated and unmethylated ZBTB33 binding sites located exactly at the peak centers (bp around peak centers), using the Fisher’s exact test.

### Peak annotation in the genomic location context of ZBTB33 binding sites

To confirm the genomic location of ZBTB33 binding sites, we utilized the *annotatePeaks* function from the HOMER package [62], where the nearest transcription start site is assigned for each peak. The annotations include, but are not limited to, promoters (defined from −1 kb to +100 bp), transcription termination sites (defined from −100 bp to +1 kb), coding DNA sequence exons, 5’ UTR exons, 3’ UTR exons, introns, and intergenic regions.

### Gene ontology analysis of ZBTB33 target genes

The genomic regions enrichment of annotations tool (GREAT) [26] was used to predict functions of ZBTB33 target genes. Because GREAT is restricted to the analysis of sequencing experiments performed in the hg19 assembly, we converted our ZBTB33 ChIP-seq peaks coordinates from hg38 to hg19 using the *LiftOver* program hosted at the UCSC genome browser website (http://genome.ucsc.edu/cgi-bin/hgLiftOver, Supplementary Table 13).

### Chromatin landscapes and co-occupancy of other transcription factors surrounding un/methylated ZBTB33 binding sites

To characterize histone modification profiles and RNA polymerase II activities surrounding un/methylated ZBTB33 binding sites, we downloaded K562 and GM12878 ChIP-seq datasets for active histone marks (H3K4me1, H3K4me3 and H3K27ac), repressive histone marks (H3K9me3 and H3K27me3) and RNA polymerase II (POLR2A, POLR2AphosphoS2, and POLR2AphosphoS5) from the ENCODE consortium and GEO datasets (Supplementary Tables 14 and 15). In addition, FAIRE-seq, DNase-seq (DUKE and UW) and ATAC-seq datasets were applied as chromatin state indicators (Supplementary Tables 14 and 15). In order to compare H3K4me1 enrichment surrounding the CpGs found in methylated ZBTB33 binding sites with genome-wide all methylated CpGs as well as the CpGs found in unmethylated ZBTB33 binding sites, H3K4me1 read intensities, normalized by sequencing depth, were accordingly extracted in a 5bp window size surrounding them and further underwent the inverse hyperbolic sine transformation.

The co-occupancy of transcription factors associated with un/methylated ZBTB33 binding sites was calculated using 221 and 132 ChIP-seq raw datasets in K562 and GM12878 respectively, downloaded from the ENCODE consortium website (Supplementary Table 16). After a read base quality check and read trimming when necessary using Trimmomatic [63], ChIP-seq reads were aligned to the human genome reference hg38 using STAR with the following parameters: --outFilterMultimapNmax 1, -- outFilterMatchNminOverLread 0.80 and blocked splice function [50]. We segregated these ChIP-seq assays into two groups according to the number of unique map reads. ChIP-seq assays with more than 30 million unique map reads were considered high quality, and ChIP-seq assays with less than 30 million unique map reads were considered lower quality.

Heatmap files of read intensities normalized by sequencing depth in the ±5kb regions surrounding ZBTB33 binding sites were generated with the *annotatePeaks* module from the HOMER package using a resolution of 25 bp, to portray the chromatin and epigenetic landscapes. Similar strategy as aforementioned was employed to profile the co-occupancy of transcription factors surrounding ZBTB33 un/methylated binding sites in a ±500 bp window size with a resolution of 5 bp. In addition, the fold enrichment of each transcription factor at ZBTB33 binding sites was calculated as the ratio of the average read intensity in the ±100 bp region surrounding the peak center to the average read intensity in a 200 bp window located 300 bp away from the peak center. Hierarchical clustering and heatmap plots were processed using the heatmap.2 function from the R gplots library (https://cran.r-project.org/web/packages/gplots/index.html).

### Integration of K562 un/methylated ZBTB33 binding sites with ChromHMM and ENCODE TFBS clusters

ENCODE TFBS clusters v3 and ChromHMM files were downloaded from the UCSC genome browser (https://genome.ucsc.edu, Supplementary Table 17). Since ENCODE TFBS clusters and Broad ChromHMM are restricted to the hg19 genome assembly, we used the *LiftOver* program from UCSC (http://genome.ucsc.edu/cgi-bin/hgLiftOver) to convert our ZBTB33 ChIP-seq peaks from the hg38 to the hg19 release (Supplementary Table 13).

We defined a region of ±100 bp surrounding each ZBTB33 un/methylated peak summit as its peak region. ZBTB33 peak regions were overlapped with the ENCODE TFBS clusters and domains defined by the ChromHMM database using the *IntersectBed* function from the BEDTools package [64]. Regarding the comparison with TFBS clusters, we counted the number of overlaps between ZBTB33 peak regions and TFBS hits for each transcription factor available in the TFBS cluster database. Then, this count (*Z*) was normalized as following: given *Y* as the total number of ZBTB33 peak regions and as the total number of experiments (ChIP-seq), then 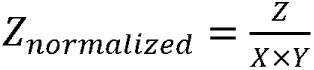.

## List of abbreviations

WGBS: whole genome bisulfite sequencing
5-aza-dC: 5-aza-2’-deoxycytidine
RRBS: reduced representation bisulfite sequencing
GO: gene ontology
GREAT: the genomic regions enrichment of annotation tool
FAIRE-seq: formaldehyde-assisted isolation of regulatory element sequencing
DNase-seq: DNase I hypersensitive site sequencing
ATAC-seq: assay for transposase-accessible chromatin sequencing
IDR: irreproducible discovery rate
PSWM: position-specific weight matrix

## Declarations

### Ethics approval and consent to participate

Not applicable

### Consent for publication

Not applicable

### Availability of data and materials

The datasets analysed during the current study are available in ENCODE consortium.

### Competing interests

The authors declare that they have no competing interests.

### Funding

Work in the T.B. laboratory is supported by the National Research Foundation, the Singapore Ministry of Education under its Centres of Excellence initiative, the National Medical Research Council of Singapore grant number NMRC/BNIG/2035/2015, the Singapore Ministry of Education Academic Research Fund Tier 3, grant number MOE2014-T3-1-006, and the Institut Français à Singapour (Merlion Project grant number 6.10.14). S.J. was supported by grants from the National Research Foundation Singapore and the Singapore Ministry of Education under its Research Centers of Excellence initiative to the Cancer Science Institute of Singapore (R-713-006-014-271), National Medical Research Council (NMRC CBRG- NIG BNIG11nov001), Ministry of Education Academic Research Fund (MOE AcRF Tier 1 T1-2012 Oct - 04 and T1-2016 Apr -01) and by the RNA Biology Center at CSI Singapore, NUS, from funding by the Singapore Ministry of Education’s Tier 3 grants, grant number MOE2014-T3-1-006.

### Authors’ contributions

T.B. made the original observations; Q.X.X.L. performed all computational analysis; Q.X.X.L., K.R., S.J. and T.B. analyzed and interpreted the data; Q.X.X.L. and T.B. wrote the paper; T.B. directed the study.

## Acknowledgements

The authors thank Ong Chin Tong, Daniel G. Tenen and Denis Thieffry for useful discussions and comments during the preparation of this manuscript, as well as Celestina Chin Ai Qi for her proofreading.

