## Supplementary Figures for "ZBTB33 (Kaiso) methylated binding sites are associated with primed heterochromatin"

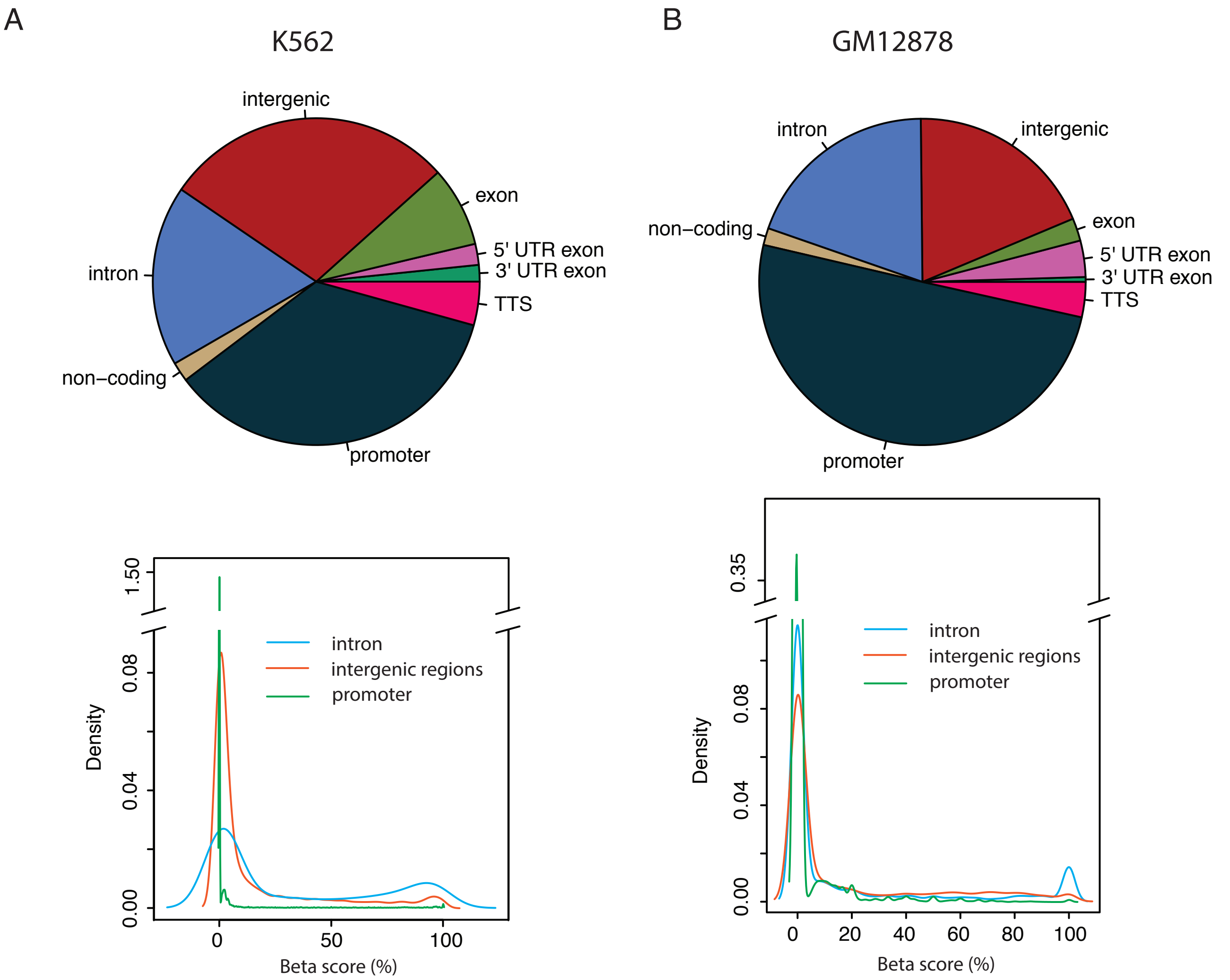


**Supplementary Figure 1. Peak annotations and DNA methylation profiling in previous ZBTB33 ChIP-seq data in K562 and GM12878.** Peak regions identified using IDR pipeline from the previous ZBTB33 ChIP-seq datasets were annotated in K562 and GM12878 (**A** and **B** respectively). The DNA methylation levels in peaks located in the promoters, intron and intergenic regions were profiled as the density plots (green, blue and orange lines respectively).

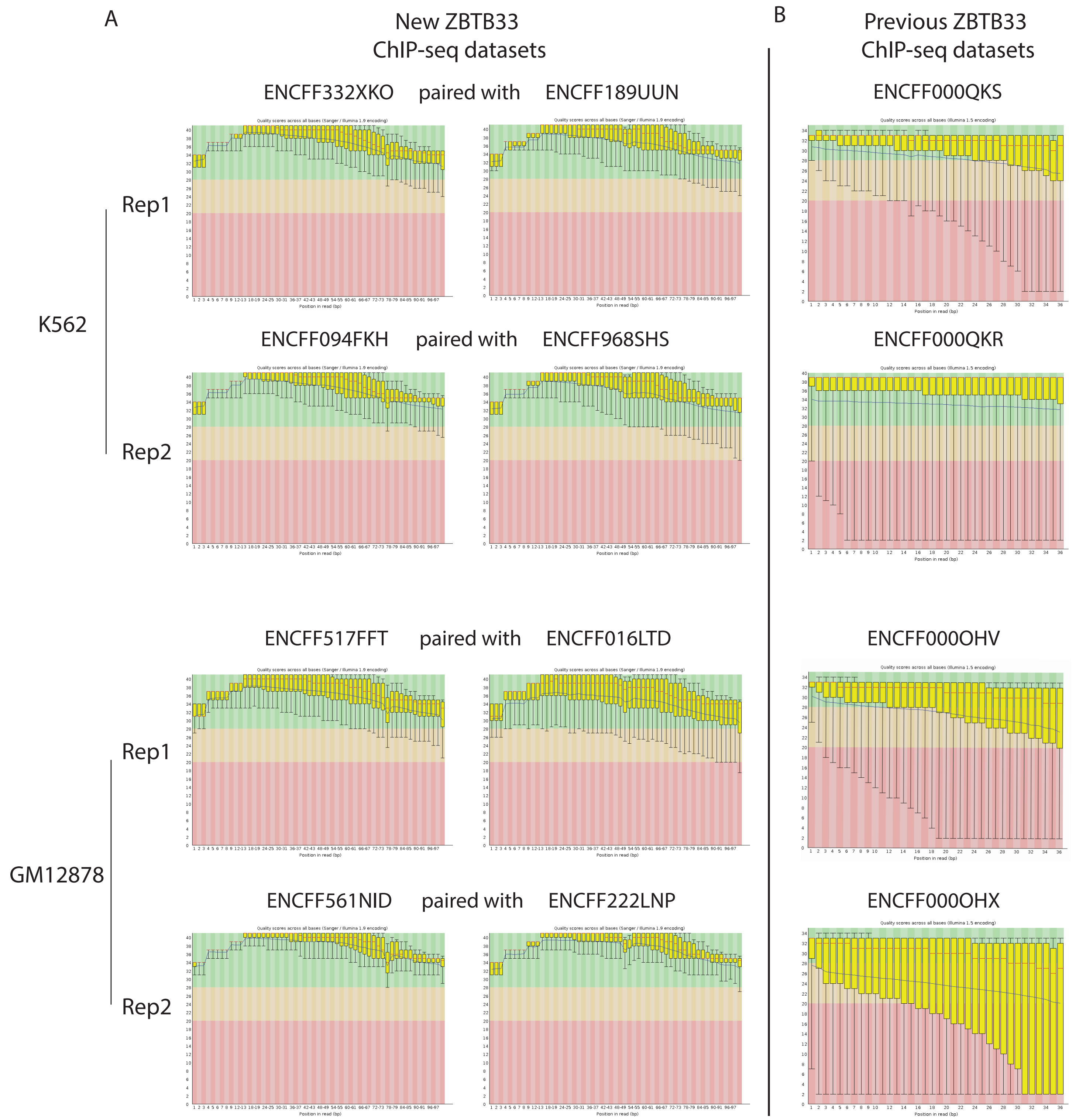


**Supplementary Figure 2. Read quality in the latest and previous ChIP-seq datasets.** Read quality was analyzed by fastQC for the latest **(A)** and previous **(B)** ZBTB33 ChIP-seq datasets in K562 (upper panel) and GM12878 (lower panel).

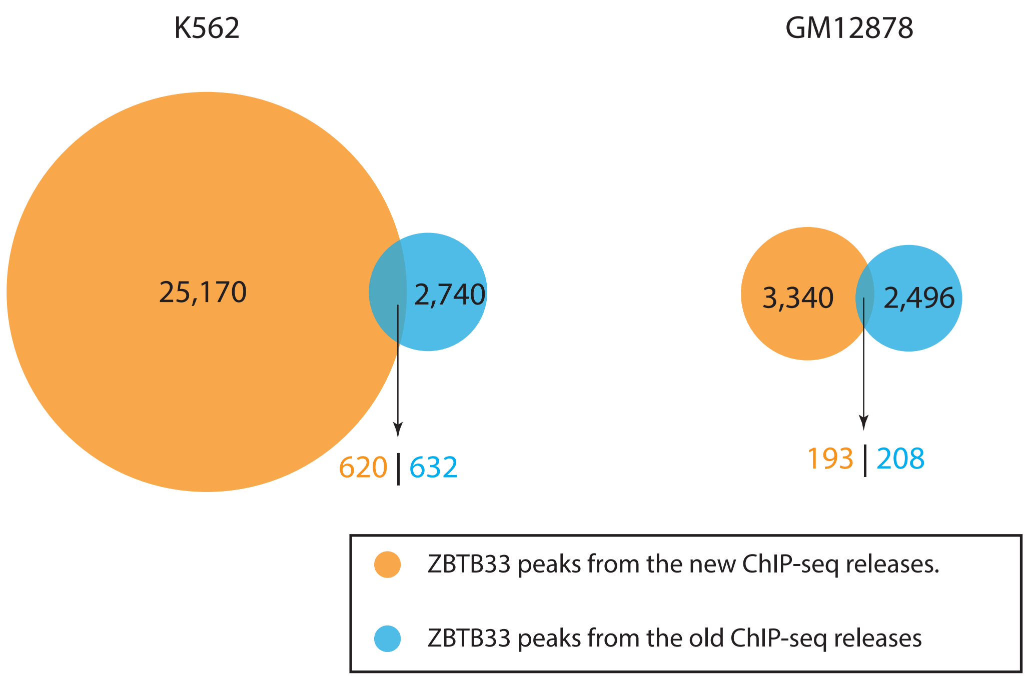


**Supplementary Figure 3. Comparison of ZBTB33 binding sites from two ENCODE ChIPseq releases.** Venn Diagrams represent the overlap of ZBTB33 IDR peaks from the new ENCODE ChIP-seq dataset releases (orange) with the IDR peaks called from previous ChIP-seq versions (blue). In K562, only 620 out of total 25,170 peaks from the latest ChIP-seq release overlap with the old ZBTB33 ChIP-seq peak version, while 632 out of 2,740 ZBTB33 ChIP-seq peaks called from the old version are common with the peaks from the latest ZBTB33 ChIP-seq experiment. In GM12878, only 193 out of 3,340 peaks from the latest ChIP-seq release overlap with the old ZBTB33 ChIP-seq peak version, while 208 out of 2,496 ZBTB33 ChIP-seq peaks called from the old version are common with peaks from the latest ZBTB33 ChIP-seq peaks.

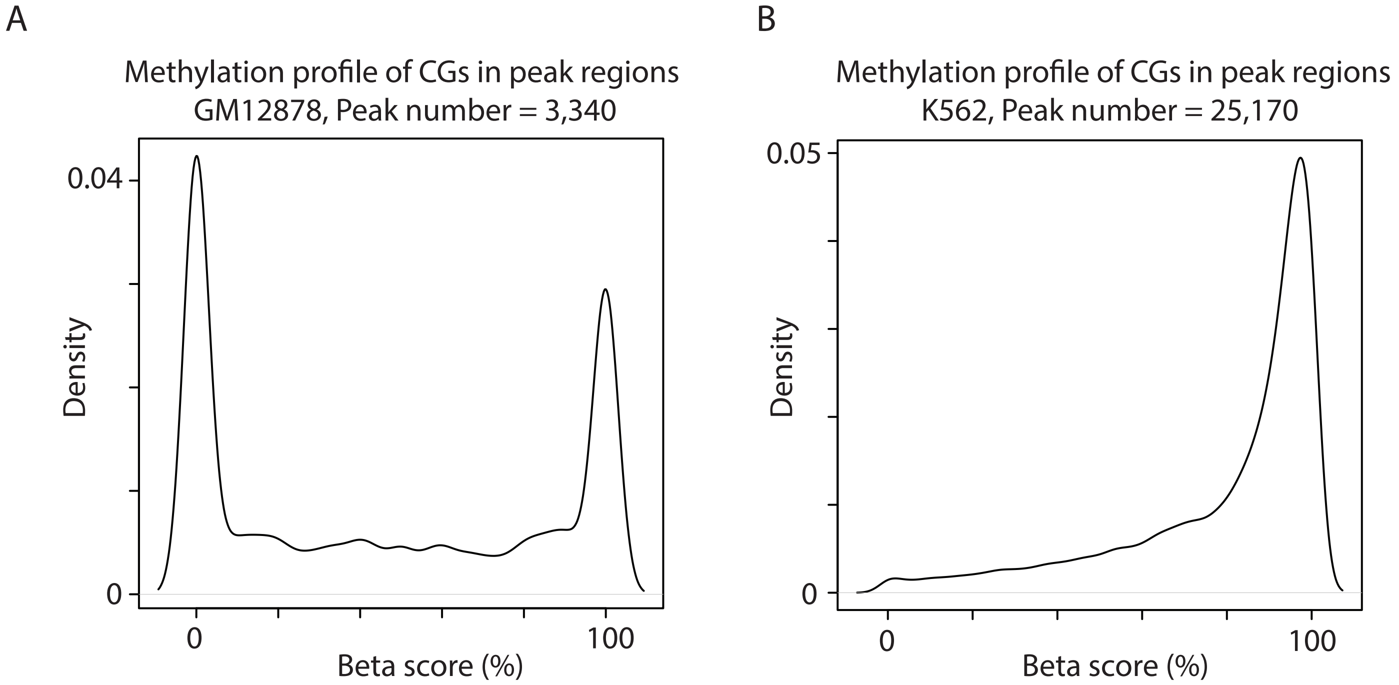


**Supplementary Figure 4. Methylation profiles of ZBTB33 peak regions in GM12878 and K562.** Densities of CpG beta scores (methylation levels) in ZBTB33 peak regions were profiled in GM12878 (A) and K562 (B). Here, only CpGs with at least 5 reads $\times$ 1 replicate and 5 reads $\times$ 2 replicates sequencing coverage were plotted in GM12878 and K562 respectively.

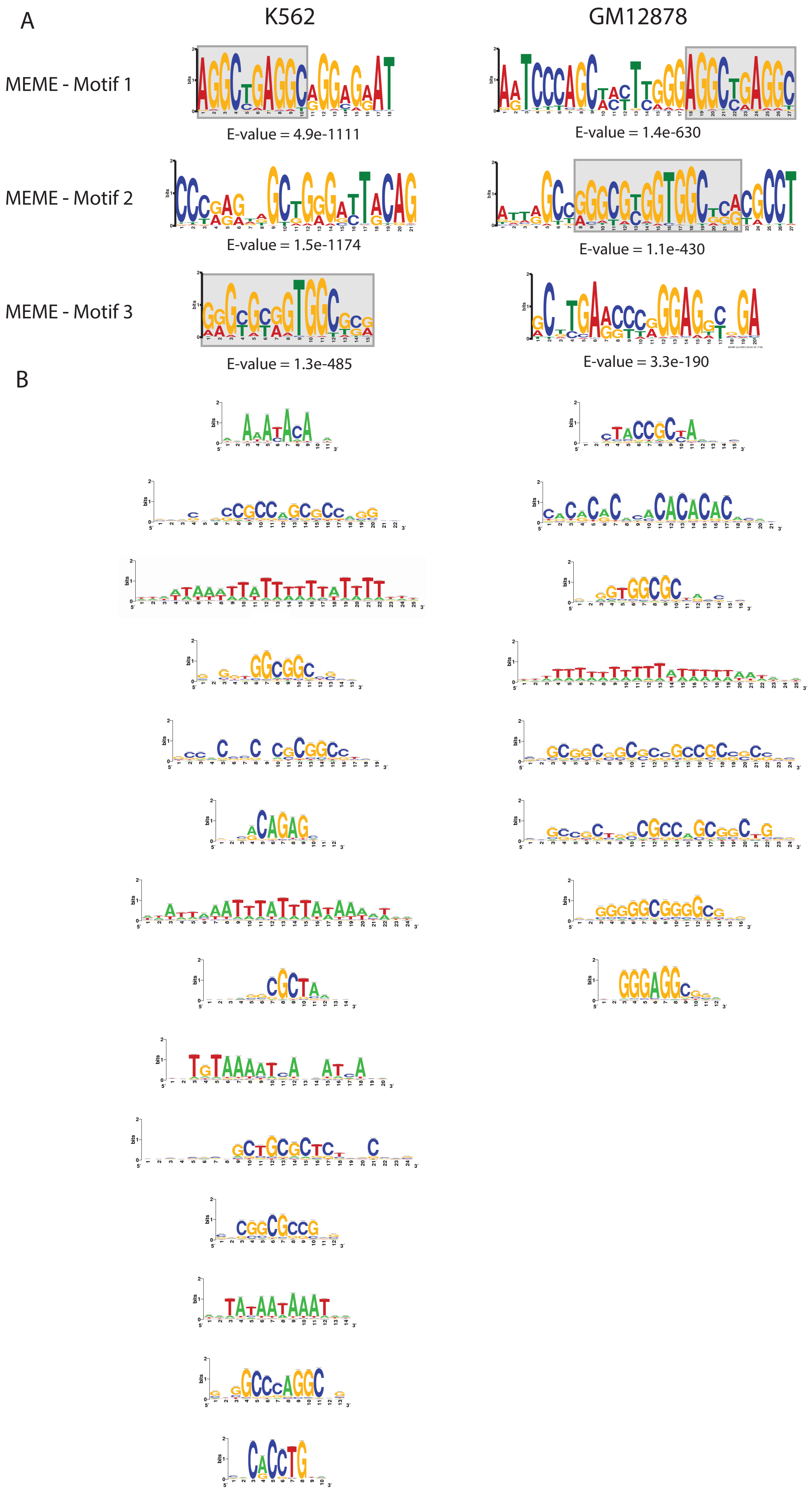


**Supplementary Figure 5. Motif de novo discovery by MEME and peak-motifs module in RSAT.** Motifs were searched de novo in 25,170 and 3,340 ZBTB33 peaks in K562 and GM12878 respectively using MEME-ChIP **(A)** and peak-motifs module in RSAT **(B)**.

Supplementary Figure 5
Lin et. al

A

K562

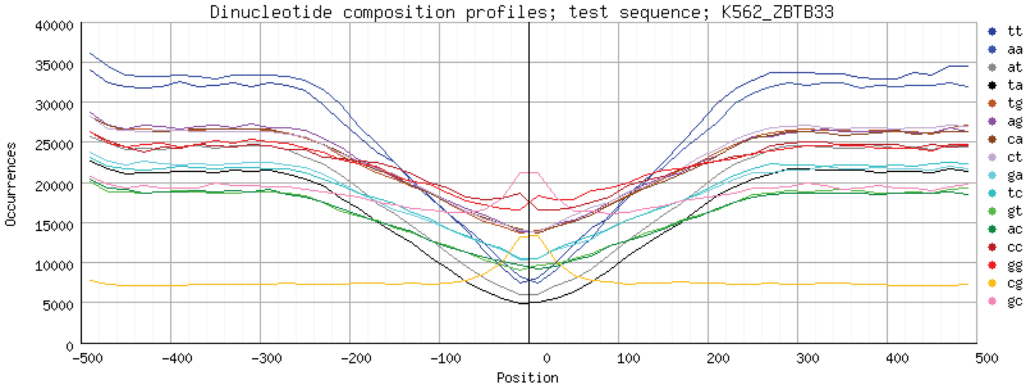


B

GM12878

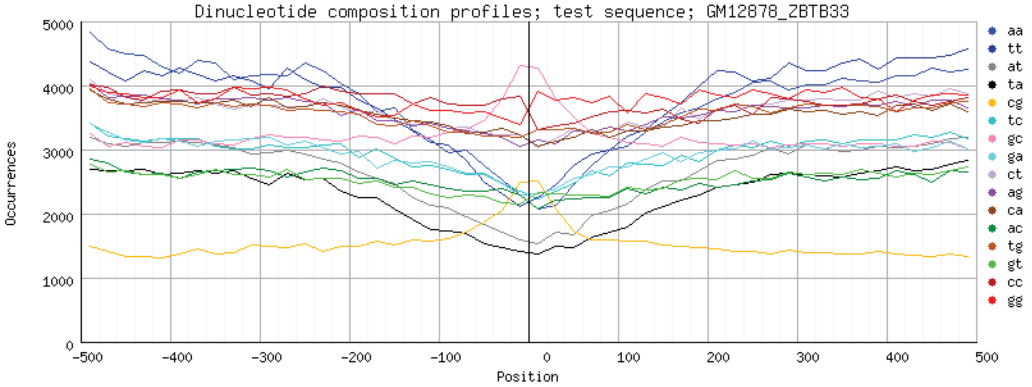


**Supplementary Figure 6: Dinucleotide composition profiles surrounding ZBTB33 peak summits in K562 and GM12878.** The dinucleotide compositions are profiled surrounding all identified ZBTB33 peak summits in K562 **(A)** and GM12878 **(B)**.

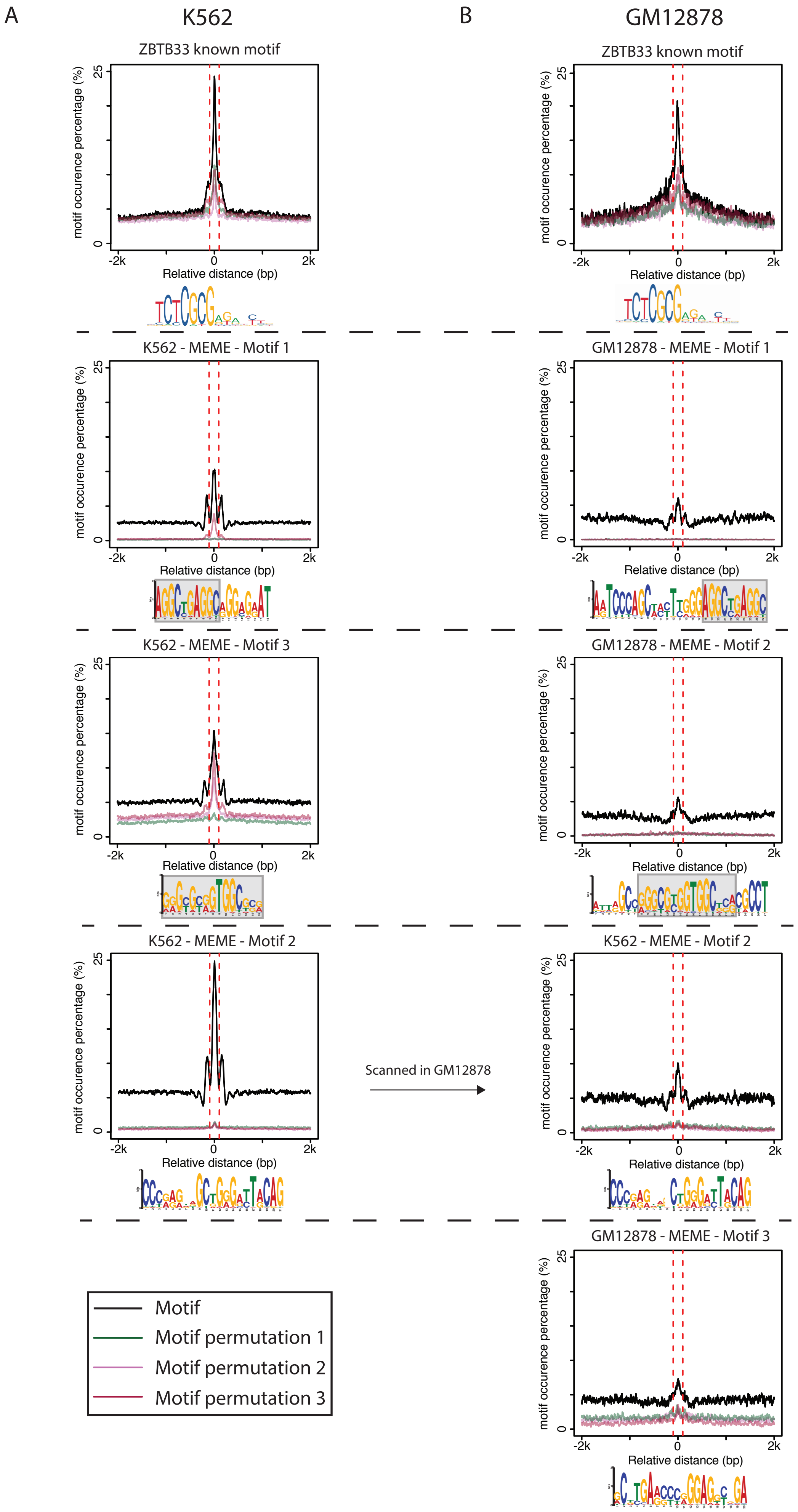


**Supplementary Figure 7: Comparison of motif occurrences between the known ZBTB33 motif and MEME motifs in K562 and GM12878.** The motif occurrences of the known ZBTB33 motif and ones de novo discovered by MEME in K562 and GM12878 were scanned surrounding peaks in the respective cell lines. In particular, the motif 2 found by MEME in K562 was scanned in GM12878 ZBTB33 peaks to test the conservation of the motif in GM12878.

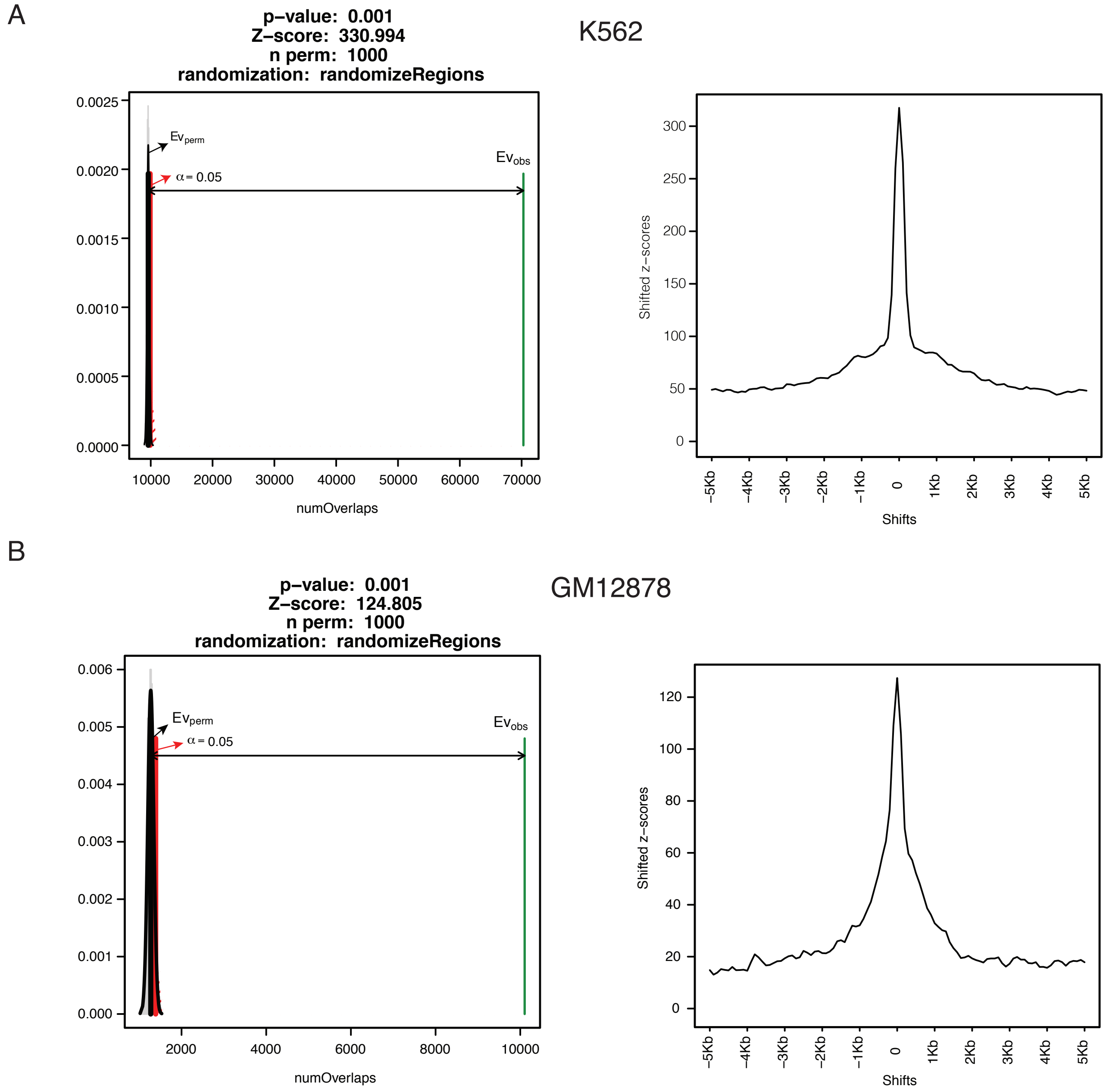


**Supplementary Figure 8: Motif enrichment analysis by regioneR in K562 and GM12878.** The original overlap between genome-wide known ZBTB33 motif (TCTCGCGAGA) and ZBTB33 peak regions (±100 bp around peak summits) is compared with randomized overlap distribution which is generated by 1000 permutation tests (left side). In each permutation test, the overlap between the randomized regions and the genome-wide potential ZBTB33 binding sites was calculated (grey area). The black line represents the mean of overlaps in the permutation tests, the green line shows the original overlap between genome-wide ZBTB33 motif and all identified ZBTB33 peaks, and the red line denotes the significant limit. A local profile of z-scores around ZBTB33 peaks in the window size of 10kb was drawn to show the dependency of the original overlap association on the exact position surrounding ZBTB33 peak centers (right side).

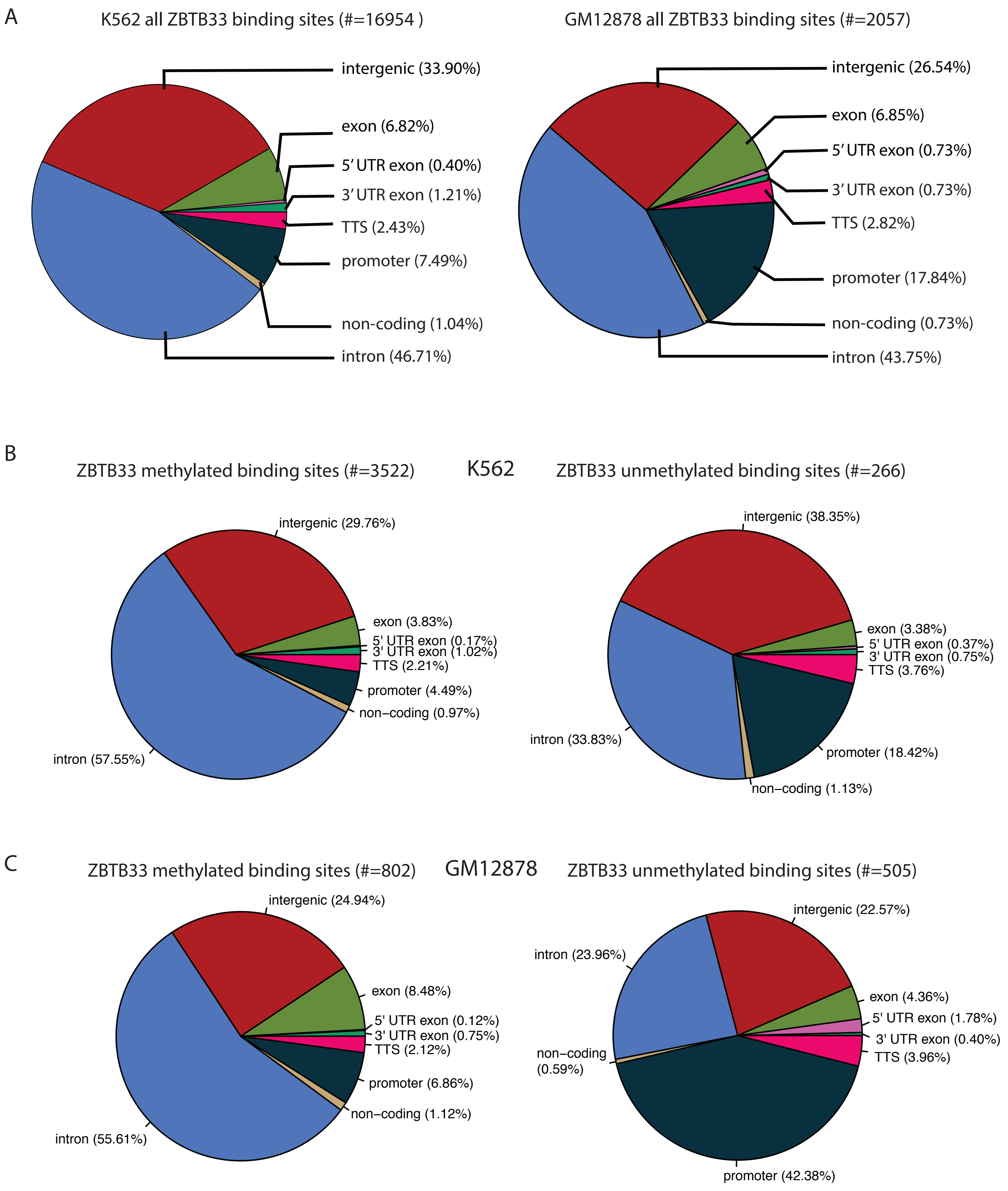


**Supplementary Figure 9. Genomic locations of ZBTB33 binding sites.** All ZBTB33 binding sites (A) as well as methylated and unmethylated binding sites (K562 in B and GM12878 in C) were annotated according to their genomic locations using the annotatePeaks function from the HOMER package.

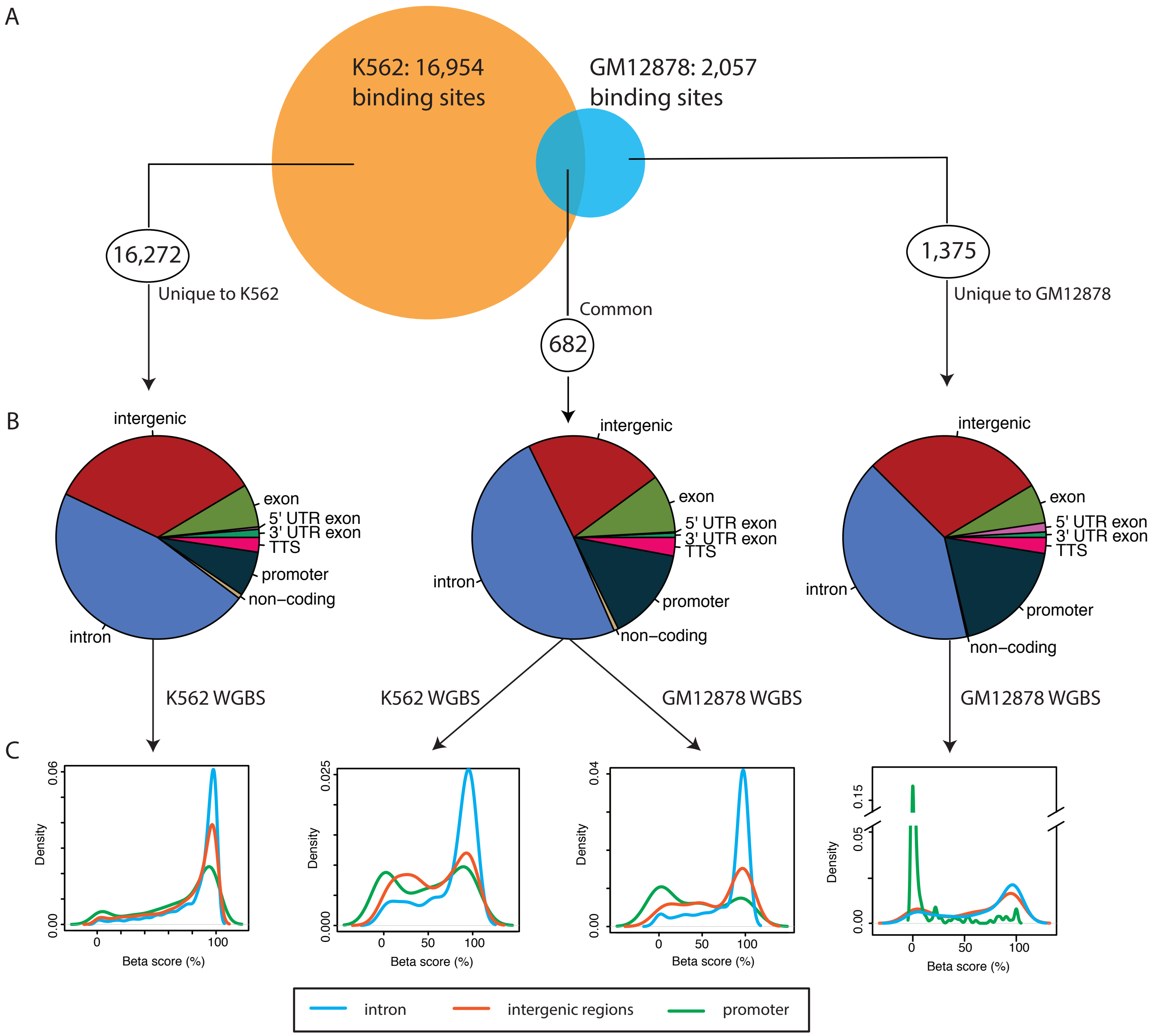


**Supplementary Figure 10. Comparison of ZBTB33 binding sites from the newly deposited ChIP-seq in K562 and GM12878. A)** Venn diagram shows the overlap of the identified 16,954 and 2,057 ZBTB33 binding sites in K562 and GM12878 respectively, only 682 ZBTB33 binding sites are common in both cell lines. **B)** Pie charts represent the genomic locations of 682 shared ZBTB33 binding sites, 16,272 binding sites unique to K562 and 1,375 binding sites unique to GM12878. **C)** Density plots display the methylation levels of shared ZBTB33 binding sites (middle two), ZBTB33 binding sites unique to K562 (extreme left) and ZBTB33 binding sites unique to GM12878 (extreme right).

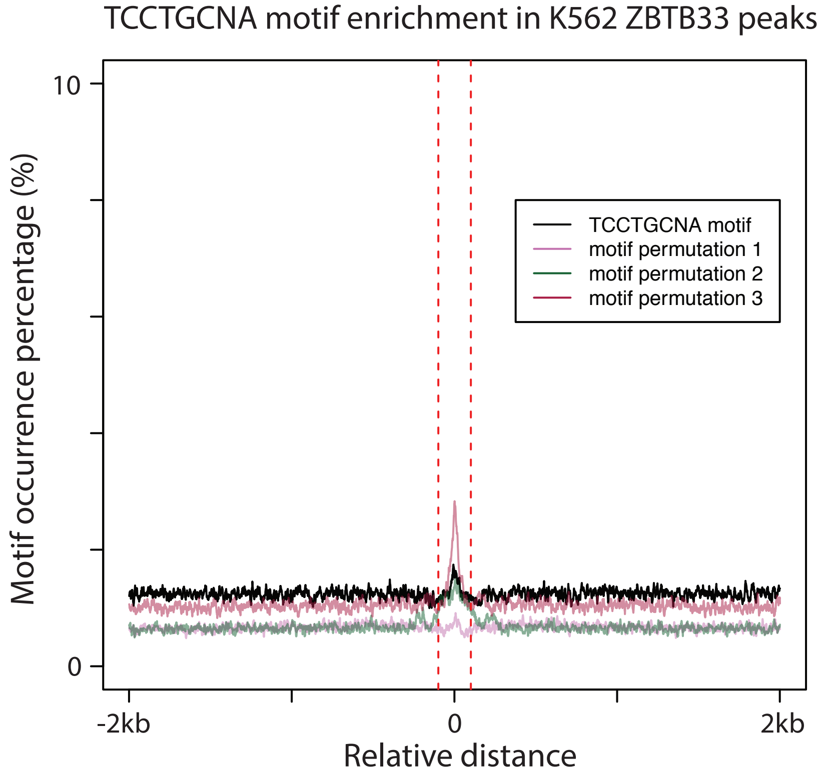


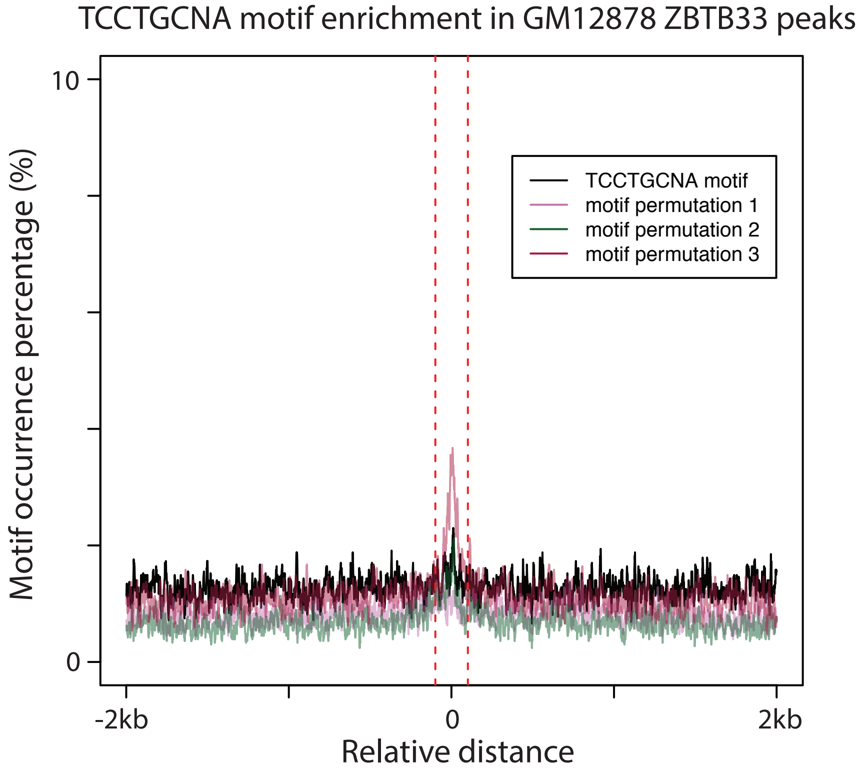


**Supplementary Figure 11. TCCTGCNA motif analysis surrounding ZBTB33 peaks.** Occurrences of the TCCTGCNA motif within the +/- 2 kb regions surrounding all 25,170 ZBTB33 ChIP-seq peaks in K562 (top) and all 3,340 ZBTB33 ChIP-seq peaks in GM12878 (bottom) were computed by matrix-scan from RSAT (black line). The significance of the motif enrichment is assessed by comparing the TCCTGCNA motif enrichment with 3 permutated motifs derived from the TCCTGCNA motif matrix (pink, green and red lines).

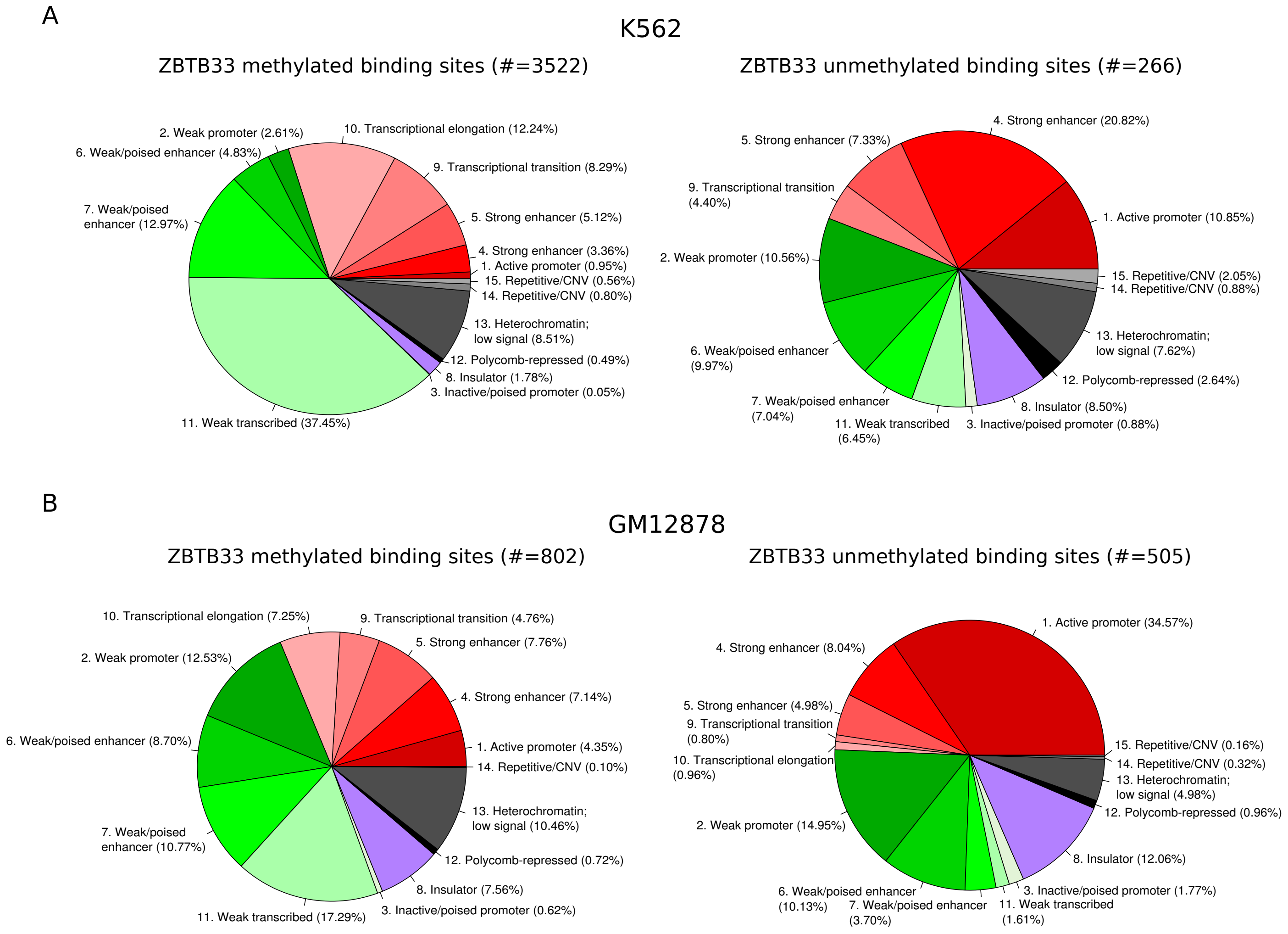


**Supplementary Figure 12. Chromatin functional states of methylated and unmethylated ZBTB33 binding sites.** Pie charts describe the functional annotations of methylated (left) and un-methylated (right) ZBTB33 binding sites predicted by ChromHMM database in K562 (A) and GM12878 (B).

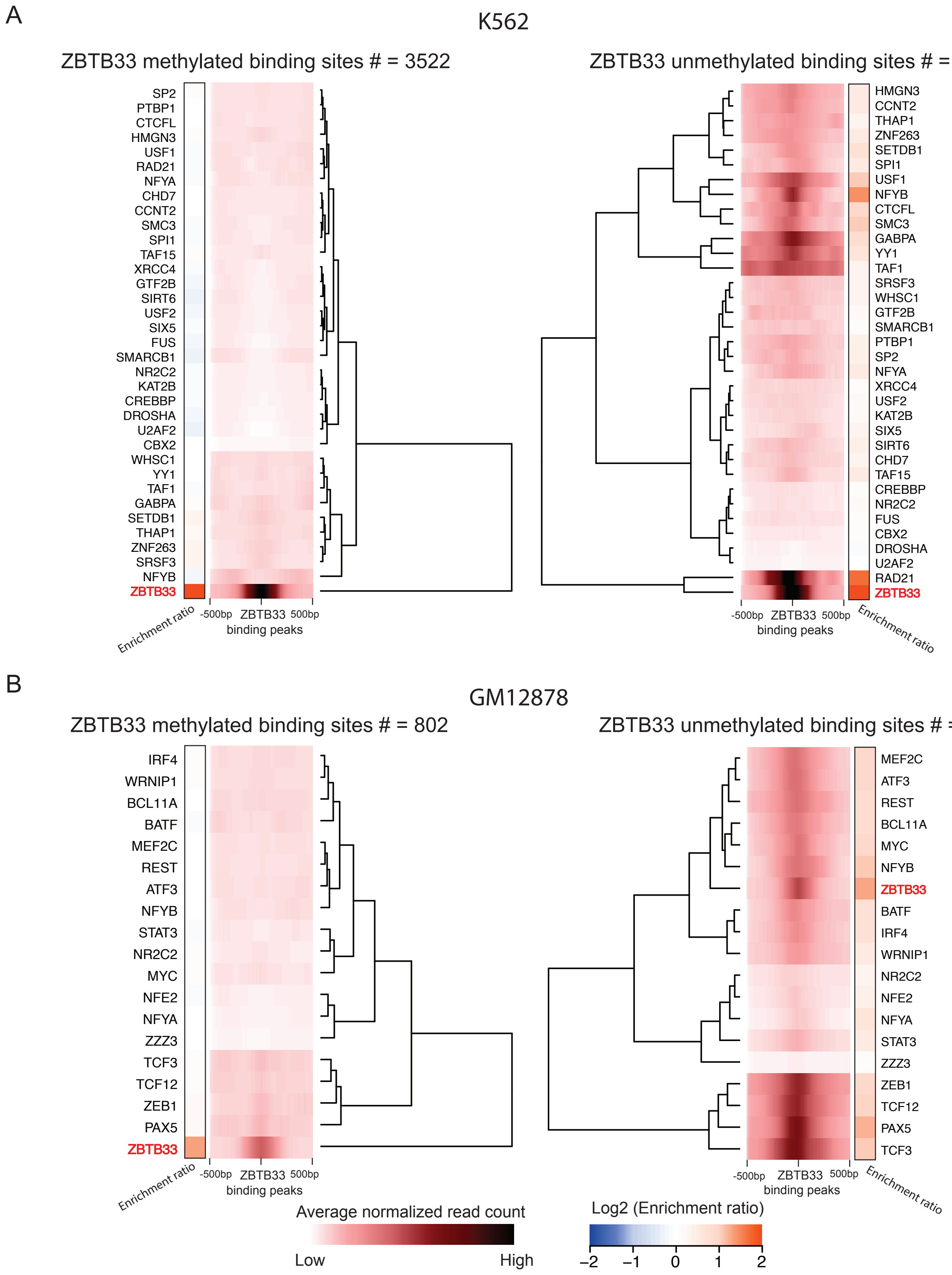


**Supplementary Figure 13. Co-occupancy between transcription factors and ZBTB33 (un)methylated binding sites.** Heatmaps describe the enrichment (average normalized read count) of 34 and 18 transcription factors in K562 (A) and GM12878 (B) respectively from ENCODE datasets (ChIP-seq experiments containing less than 30 million uniquely mapped reads) within +/- 500 bp regions surrounding methylated (left) and unmethylated (right) ZBTB33 binding sites. Here, each row represents a sequencing experiment from ENCODE where read intensities for all ZBTB33 peak regions were averaged, while each column represents relative distance to ZBTB33 peak center. Furthermore, the fold enrichment of the average normalized read counts in the peak regions (+/- 100 bp around peak centers) compared to the flanking regions (200 bp windows located 300 bp away from the peak centers) for each TF is shown in a heatmap. Orange color represents the average normalized read count in the peak regions is more than in the flanking regions, while the blue color denotes the opposite.

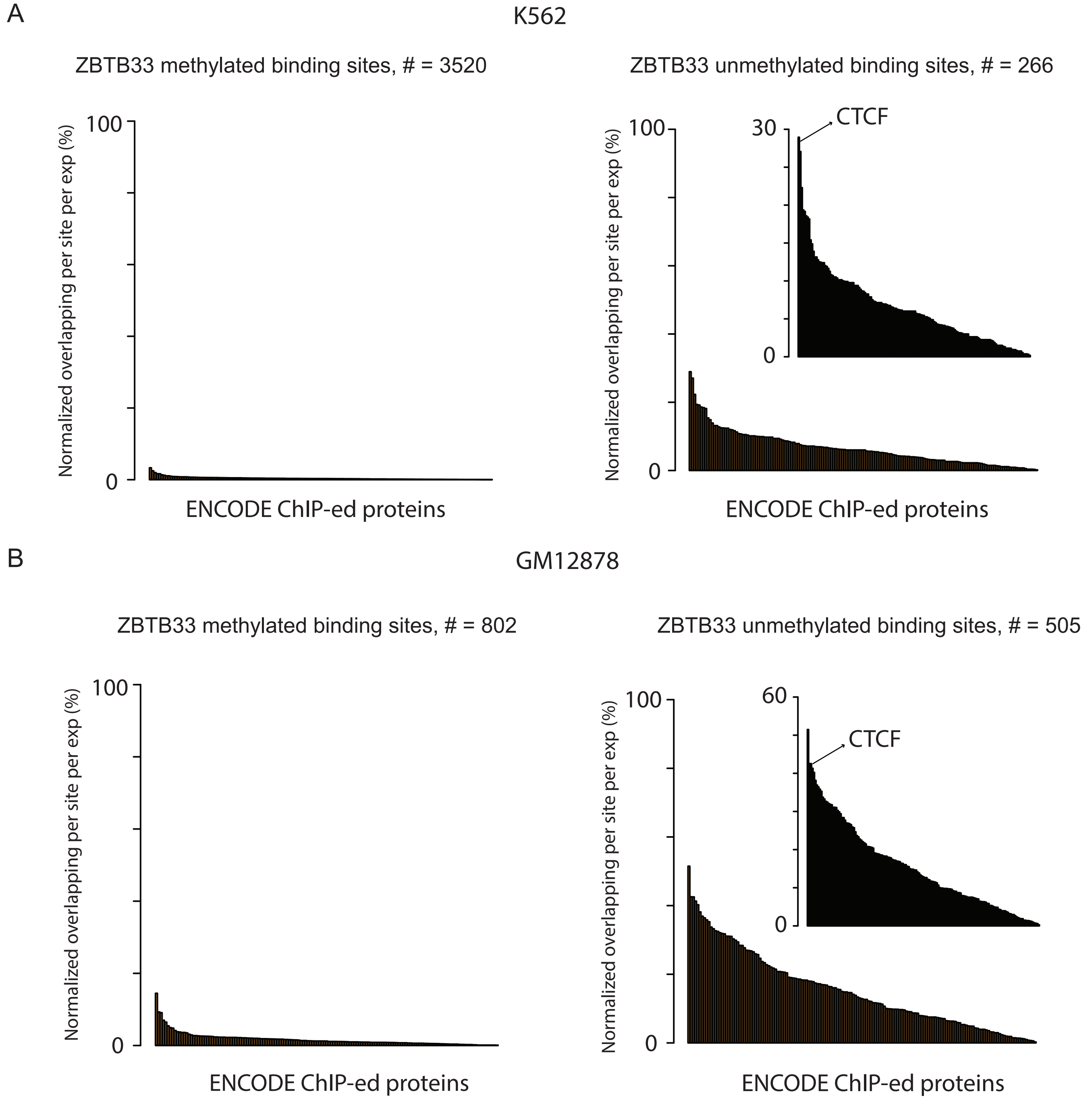


**Supplementary Figure 14. Overlap between ZBTB33 binding sites and ENCODE TFBS clusters.** Methylated (left) and unmethylated (right) ZBTB33 binding sites in K562 (A) and GM12878 (B) were overlapped with the ENCODE TFBS clusters. The number of overlaps between ZBTB33 peak regions and TFBS sites for each transcription factor available in the TFBS cluster database was counted. Then, this count (Z) was normalized as following: given X as the total number of ZBTB33 peak regions and Y as the total number of experiments (ChIP-seq), then $Z_{normalized}=Z/(X \times Y)$.

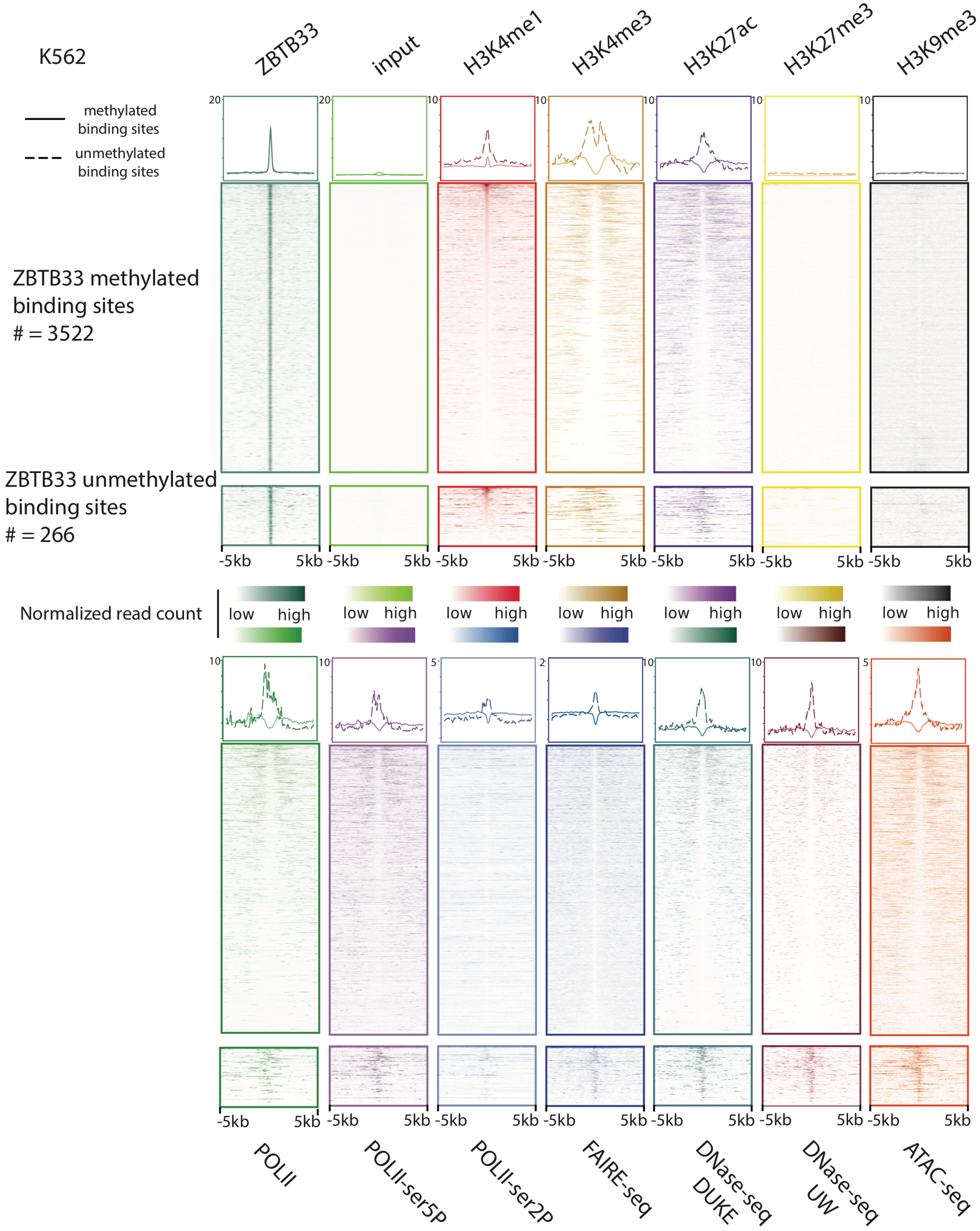


**Supplementary Figure 15. Chromatin landscape surrounding methylated and unmethylated ZBTB33 binding sites in K562.** Heatmaps show the enrichment (normalized read count) of histone marks, RNA polymerase II and datasets indicating chromatin states within +/- 5 kb regions surrounding methylated and unmethylated ZBTB33 binding sites (from 3rd panel to 14th panel). The average read intensities were shown on the top of the heatmaps (solidline: methylated ZBTB33 binding sites; dash line: unmethylated ZBTB33 binding sites). Here each row represents a ZBTB33 binding site region in the descending order of H3K4me1 read intensity (third panel), and each column represents the relative distance from the ZBTB33 ChIPseq peak center.

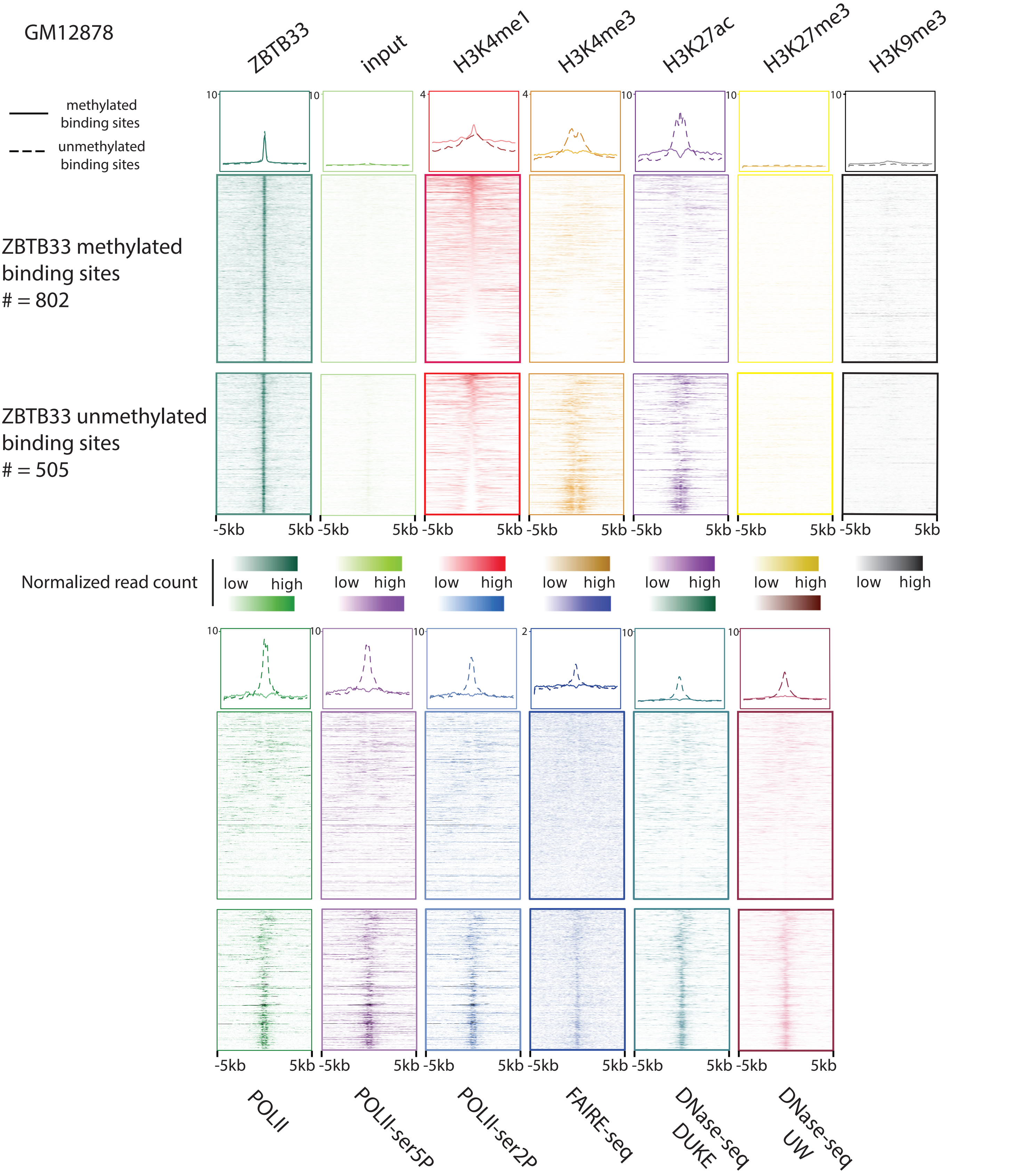


**Supplementary Figure 16. Chromatin landscape surrounding methylated and unmethylated ZBTB33 binding sites in GM12878.** Heatmaps show the enrichment (normalized read count) of histone marks, RNA polymerase II and datasets indicating chromatin states within +/- 5 kb regions surrounding methylated and unmethylated ZBTB33 binding sites (from 3rd panel to 13th panel). The average read intensities were shown on the top of the heatmaps (solidline: methylated ZBTB33 binding sites; dash line: unmethylated ZBTB33 binding sites). Here each row represents a ZBTB33 binding site region in the descending order of H3K4me1 read intensity (third panel), and each column represents the relative distance from the ZBTB33 ChIPseq peak center.

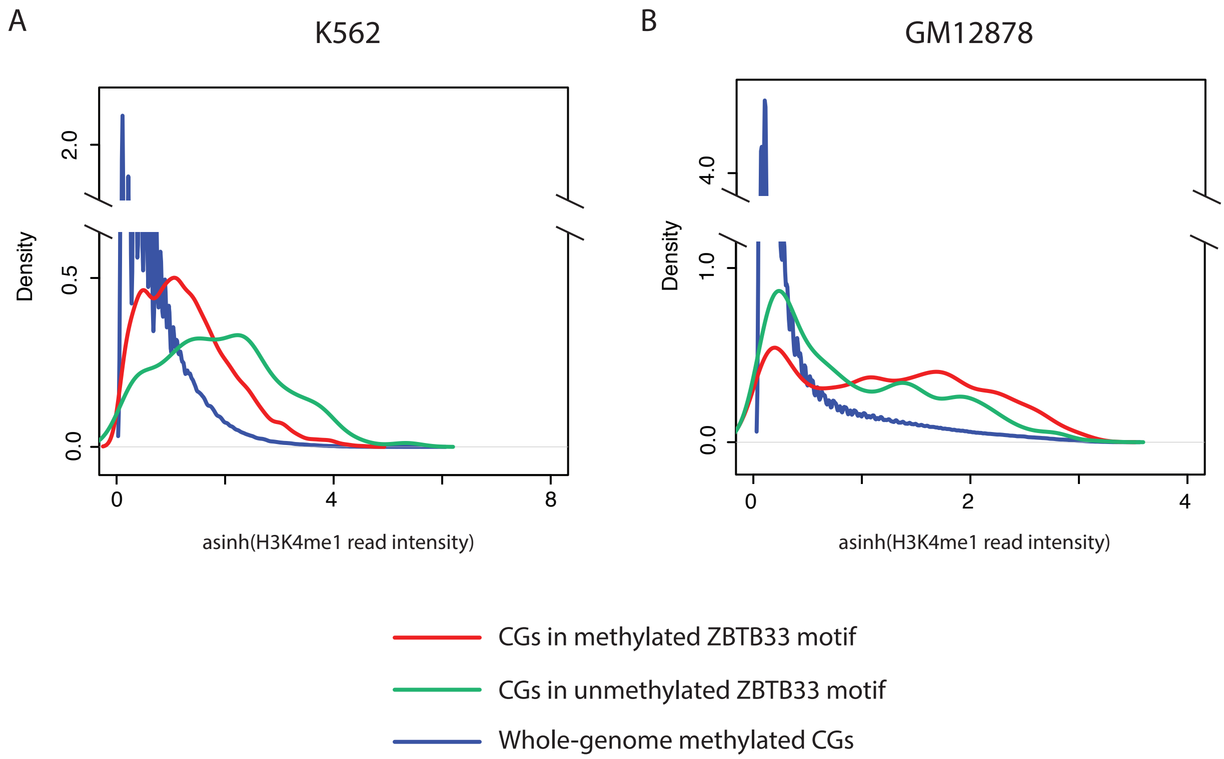


**Suplementary Figure 17: H3K4me1 read intensities around all methylated CpGs, CpGs in methylated ZBTB33 motif and CpGs in unmethylated ZBTB33 motif in K562 and GM12878.** The density plots show the distributions of the inverse hyperbolic sine transformed H3K4me1 read intensities around all methylated CpGs, CpGs in methylated ZBTB33 motifs and CpGs in unmethylated ZBTB33 motifs (blue, red and green lines respectively) in K562 **(A)** and GM12878 **(B)**.
