## Supplementary Tables for "ZBTB33 (Kaiso) methylated binding sites are associated with primed heterochromatin"

### **Supplementary Table 1.** ZBTB33 ChIP-seq datasets used in previous study [1]

| Cell line | Accession | Antibody | Released date | Uniquely mapped reads (rep1; rep2) | Run type [2] | self-pseudoreplicates from replicate | self-pseudoreplicates from replicate | Original replicates [3] | Pooled pseudoreplicate [4] | Final peak (conservative IDR peak) | Input control |
|---|---|---|---|---|---|---|---|---|---|---|---|
| GM12878 | ENCSR000BHC | ENCAB000AML | 18/07/2011 | 13,936,377; 12,919,213 | SE36nt | 2848 | 2940 | 2496 | 2126 | 2496 | ENCSR000BGH |
| K562 | ENCSR000BKF | | 18/07/2011 | 25,117,058; 14,890,947 | SE36nt | 8795 | 6406 | 2740 | 6345 | 2740 | ENCSR000BLJ |

Note: irreproducibility Discovery Rate (IDR) pipeline was used for reproducibility test and peak calling. In ENCODE, the recommended rescue and self-consistency ratios are less than 2. Rescue ratio in G12M878: 2496/2126<2; self-consistency ratio in GM12878: 2940/2848<2; rescue ratio in K562: 6345/2740>2; self-consistency ratio in K562: 8795/6406<2.

[1] Blattler, A. et al. ZBTB33 binds unmethylated regions of the genome associated with actively expressed genes. Epigenetics Chromatin 6, 13 (2013).

[2] SE - single end

[3] Peaks called from the original replicates; IDR thresholds < 0.05

[4] Peaks called from the pseudo-replicates; IDR thresholds < 0.01

### **Supplementary Table 2.** Comparison of new and previous ZBTB33 ChIP-seq

| | Latest ZBTB33 ChIP-seq | | Previous ZBTB33 ChIP-seq | |
|---|---|---|---|---|
| | K562 (Snyder) | GM12878 (Snyder) | K562 (Myers) | GM12878 (Myers) |
| Antibody primary test (immunoblot analysis) | compliant | | exempt from standards (not fulfilled the ENCODE Consortium's antibody standards but absolved by an ENCODE Antibody Review Committee) or not submitted to review by lab | |
| Antibody secondary test (mass spectrometry) | compliant. Kaiso unique peptide count is 6. Kaiso was not found in control IP | | compliant in one MS assay but not compliant in another MS assay. In compliant assay, Kaiso unique peptide count is 3 and fold enrichment is 4.53 | |
| ChIP-seq read quality | Good | Good | Concerning | Concerning |
| ChIP-seq read size[1] | compliant (Paired-end 100nt) | compliant (Paired-end 100nt) | Low read (Single end 36bp, less than recommended 50bp) | Low read (Single end 36bp, less than recommended 50bp) |
| ChIP-seq uniquely mapped reads[2] | One replicate compliant; the other shows low read (17,589,968 reads less than recommended 20M but more than acceptable limit 10M) | compliant | One replicate compliant; the other shows low read (14,890,947 reads less than recommended 20M but more than acceptable limit 10M) | both replicates show low read (both less than 20M but more than 10M) |
| Replicate consistency | compliant | compliant | Using in-house pipeline (using STAR as aligner and MACS2 as peak caller), the rescue ratio is more than 2. The recommended rescue ratio is less than 2 | compliant |
| other concerning flags in ENCODE | no other concerning flags | 1. "Mild to moderate bottlenecking". However, the concerning PCR Bottlenecking Coefficient 2 (PBC2) value is 9.96 in one replicate, slightly lower than the recommended lower limit value of 10. 2. Borderline replicate concordance. Rescue ratio values is 2.52, more than the upper limit value of 2. However, using in-house pipeline, the rescue ratio is 1.03. Such difference is due to different aligner and peak caller used. | "Mild to moderate bottlenecking". PCR Bottlenecking Coefficient 1 (PBC1) value is 0.79 in one replicate, but the recommended lower limit value is 0.9. Further, bottlenecking Coefficient 2 (PBC2) value is 4.59 in one replicate, but the recommended lower limit value is 10. | no other concerning flags |

[1] The recommended read length is more than 50bp according to ENCODE standards
[2] The recommended number of uniquely mapped reads is more than 20 million according to ENCODE standards

**Supplementary Table 3.** New ZBTB33 ChIP-seq datasets in K562 and GM12878

| Cell line | Antibody | Accession ID | Release date | Isogenic replicate | Run type [1] | Assembly | Reads | Input control |
|---|---|---|---|---|---|---|---|---|
| K562 | ENCAB292USO | ENCSR876GXA | June 23, 2016 | 1 | PE100nt | hg38 | 20,486,392 paired reads | ENCSR173USI |
| | | | | 2 | PE100nt | hg38 | 17,589,968 paired reads | |
| GM12878 | ENCAB292USO | ENCSR542FLV | November 13, 2017 | 1 | PE100nt | hg38 | 30,023,843 paired reads | ENCSR398JTO |
| | | | | 2 | PE100nt | hg38 | 35,253,588 paired reads | |

[1] PE - paired end

**Supplementary Table 4.** Peak calling of new ZBTB33 ChIP-seq datasets in K562 and GM12878 with irreproducibility discovery rate (IDR) pipeline

| Biosample | Peaks caller | Consistency analysis | | | | Final set IDR peaks after filtering blacklist [1] |
|---|---|---|---|---|---|---|
| | | Self-consistency test | | Original replicate | Pooled pseudoreplicate | |
| | | self-pseudoreplicates from replicate 1 | self-pseudoreplicates from replicate 2 | | | |
| K562 | MACS2 2.1.1 | 15,712 | 22,701 | 25,185 | 22,902 | 25,170 |
| GM12878 | MACS2 2.1.1 | 7,807 | 5,775 | 3,348 | 3,264 | 3,340 |

Note: irreproducibility Discovery Rate (IDR) pipeline was used for reproducibility test and peak calling. In ENCODE, the recommended rescue and self-consistency ratios are less than 2. Rescue ratio in K562: 25,185/22,902<2; self-consistency ratio in K562: 22,701/15,712<2; rescue ratio in GM12878: 3,348/3,264<2; self-consistency ratio in GM12878: 7,807/5,775<2.

[1] Blacklist bed file was downloaded from ENCODE and its accession is ENCSR636HFF

**Supplementary Table 5**. GenometriCorr analysis results in K562

| Item | Value |
|---|---|
| query.population | 25170 |
| reference.population | 5225554 |
| query.coverage | 5084340 |
| reference.coverage | 56019481 |
| relative.distances.ks.p.value[1] | 0 |
| relative.distances.ecdf.deviation.area | 0.05392609 |
| relative.distances.ecdf.area.correlation[2] | 0.2154702 |
| query.reference.intersection | 604741 |
| query.reference.union | 60499080 |
| jaccard.measure | 0.00999587 |
| projection.test.p.value[3] | 0 |
| projection.test.lower.tail[3] | FALSE |
| projection.test.obs.to.exp | 20.38916 |
| scaled.absolute.min.distance.sum | 3402.697 |
| reference.middles | NULL |
| relative.distances.ecdf.deviation.area.p.value[4] | <0.01 |
| scaled.absolute.min.distance.sum.p.value[4] | <0.01 |
| scaled.absolute.min.distance.sum.lower.tail[5] | TRUE |
| jaccard.measure.p.value[4] | <0.01 |
| jaccard.measure.lower.tail[6] | FALSE |

[1] Low p value suggests the locations of genome-wide ZBTB33 motif and identified ZBTB33 peaks are not independent but correlated.

[2] Positive value indicates genome-wide ZBTB33 motif and ZBTB33 peaks are closer to each other than expected.

[3] Low in projection.test.p.value and “FALSE” in projection.test.lower.tail imply significant overlap of the genome-wide ZBTB33 motif and ZBTB33 peaks.

[4] Three permutation tests give p-value<0.01, meaning that the observed the spatial relationships are significantly different from the permutation distributions.

[5] Value “TRUE” suggests the absolute distances between genome-wide ZBTB33 motif and ZBTB33 peaks are consistent and small.

[6] Value “FALSE” indicates an unexpectedly high overlap between genome-wide ZBTB33 motif and ZBTB33 peaks.

**Supplementary Table 6**. GenometriCorr analysis results in GM12878

| Item | Value |
|---|---|
| query.population | 3340 |
| reference.population | 5225554 |
| query.coverage | 674680 |
| reference.coverage | 56019481 |
| relative.distances.ks.p.value[1] | 0 |
| relative.distances.ecdf.deviation.area | 0.04763504 |
| relative.distances.ecdf.area.correlation[2] | 0.1905962 |
| query.reference.intersection | 81285 |
| query.reference.union | 56612876 |
| jaccard.measure | 0.0014358 |
| projection.test.p.value[3] | 0 |
| projection.test.lower.tail[3] | FALSE |
| projection.test.obs.to.exp | 17.32057 |
| scaled.absolute.min.distance.sum | 699.7178 |
| reference.middles | NULL |
| relative.distances.ecdf.deviation.area.p.value[4] | <0.01 |
| scaled.absolute.min.distance.sum.p.value[4] | <0.01 |
| scaled.absolute.min.distance.sum.lower.tail[5] | TRUE |
| jaccard.measure.p.value[4] | <0.01 |
| jaccard.measure.lower.tail[6] | FALSE |

[1] Low p value suggests the locations of genome-wide ZBTB33 motif and identified ZBTB33 peaks are not independent but correlated.
[2] Positive value indicates genome-wide ZBTB33 motif and ZBTB33 peaks are closer to each other than expected.
[3] Low in projection.test.p.value and "FALSE" in projection.test.lower.tail imply significant overlap of the genome-wide ZBTB33 motif and ZBTB33 peaks.
[4] Three permutation tests give p-value<0.01, meaning that the observed the spatial relationships are significantly different from the permutation distributions.
[5] Value "TRUE" suggests the absolute distances between genome-wide ZBTB33 motif and ZBTB33 peaks are consistent and small.
[6] Value "FALSE" indicates an unexpectedly high overlap between genome-wide ZBTB33 motif and ZBTB33 peaks.

**Supplementary Table 7.** (see Supplementary Table 7. ZBTB33 binding sites computed by RSAT matrix-scan in K562 and GM12878.xlsx)

**Supplementary Table 8.** genomic locations of ZBTB33 binding sites in K562

| Annotation | Number of peaks | Total size (bp) | Log2 Enrichment |
|---|---|---|---|
| Promoter | 1270 | 33835045 | 2.746 |
| Exon | 1215 | 35523244 | 2.611 |
| 5UTR | 68 | 2674233 | 2.184 |
| pseudo | 34 | 1906116 | 1.672 |
| TTS | 412 | 30225027 | 1.284 |
| ncRNA | 83 | 6143505 | 1.271 |
| 3UTR | 205 | 22600733 | 0.697 |
| Intron | 7920 | 1223137819 | 0.21 |
| Intergenic | 5747 | 1672997792 | -0.704 |
| miRNA | 0 | 84186 | -14.049 |
| snoRNA | 0 | 331 | -14.049 |
| rRNA | 0 | 16815 | -14.049 |

**Supplementary Table 9.** genomic locations of ZBTB33 binding sites in GM12878

| Annotation | Number of peaks | Total size (bp) | Log2 Enrichment |
|---|---|---|---|
| snoRNA | 1 | 331 | 12.119 |
| Promoter | 367 | 33835045 | 3.998 |
| 5UTR | 15 | 2674233 | 3.046 |
| Exon | 148 | 35523244 | 2.617 |
| pseudo | 5 | 1906116 | 1.95 |
| TTS | 58 | 30225027 | 1.499 |
| Intron | 900 | 1223137819 | 0.116 |
| 3UTR | 15 | 22600733 | -0.033 |
| Intergenic | 546 | 1672997792 | -1.057 |
| ncRNA | 2 | 6143505 | -1.061 |
| miRNA | 0 | 84186 | -11.006 |
| rRNA | 0 | 16815 | -11.006 |

**Supplementary Table 10.** (see Supplementary Table 10. Integrated analysis of ENCODE TFBS clusters with ZBTB33 methylated binding site coordinates in K562 and GM12878.xlsx)

**Supplementary Table 11.** (see Supplementary Table 11. Integrated analysis of ENCODE TFBS clusters with ZBTB33 unmethylated binding site coordinates in K562 and GM12878.xlsx)

**Supplementary Table 12.** Whole genome bisulfite sequencing (WGBS) datasets from ENCODE

| | | | | Alignment and methylation extraction | | | | | Correlation analysis and merge of replicates | |
|---|---|---|---|---|---|---|---|---|---|---|
| Experiment | Accession | Run type [1] | Reads Trimmer | Aligner and methylation extractor | | Assembly | Number of alignment with a unique best hit | Mapping efficiency | Tool | Pearson correlation coefficient |
| K562-WGBS | ENCLB742NWU | PE100nt | Trim Galore 0.4.2 | Bismark 0.16.3 | Bowtie 2 2.2.9 | hg38 | 401,243,487 | 72.70% | methylKit 0.99.3 | 0.97 |
| | ENCLB542OXH | PE125nt | | | | | 586,943,461 | 80.50% | | |
| GM12878-WGBS | ENCLB794YYH | PE125nt | | | | | 423,011,290 | 87.6% | NA | NA |

[1] PE - paired end

**Supplementary Table 13.** *LiftOver* of methylated and unmethylated ZBTB33 binding sites

| Biosample | Input | Original assembly | Original number | New assembly | New number | Success rate |
|---|---|---|---|---|---|---|
| K562 | ZBTB33 methylated binding sites | hg38 | 3522 | hg19 | 3520 | 99.94% |
| | ZBTB33 unmethylated binding sites | hg38 | 266 | hg19 | 266 | 100% |
| GM12878 | ZBTB33 methylated binding sites | hg38 | 802 | hg19 | 800 | 99.75% |
| | ZBTB33 unmethylated binding sites | hg38 | 505 | hg19 | 504 | 99.80% |

**Supplementary Table 14.** Histone mark, RNA polymerase II and chromatin state datasets in K562

| Feature | Type | Accession in ENCODE/GEO datasets | Layout | Read length | Assembly | Uniquely mapped reads number | Uniquely mapped reads percentage | Total read number |
|---|---|---|---|---|---|---|---|---|
| Active histone marks | H3K4me1 | ENCFF000VDU | Single end | 41 | hg38 | 25,985,660 | 71.70% | 35,870,715 |
| | | ENCFF000VDV | Single end | 41 | hg38 | 9,885,055 | 70.79% | |
| | H3K4me3 | ENCFF001FXG | Single end | 36 | hg38 | 15,633,931 | 83.67% | 29,732,080 |
| | | ENCFF001FXH | Single end | 36 | hg38 | 14,098,149 | 83.08% | |
| | H3K27ac | ENCFF000BXG | Single end | 36 | hg38 | 5,973,983 | 80.98% | 17,993,379 |
| | | ENCFF000BXH | Single end | 36 | hg38 | 12,019,396 | 76.97% | |
| Repressive histone marks | H3K9me3 | ENCFF001QWW | Single end | 36 | hg38 | 15,846,561 | 75.10% | 49,398,735 |
| | | ENCFF001QWX | Single end | 36 | hg38 | 33,552,174 | 72.58% | |
| | H3K27me3 | ENCFF915XIL | Single end | 36 | hg38 | 13,295,069 | 77.81% | 36,262,039 |
| | | ENCFF330YFF | Single end | 36 | hg38 | 22,966,970 | 84.07% | |
| RNA polymerase II | POLR2A | ENCFF000QDX | Single end | 36 | hg38 | 29,674,841 | 79.73% | 56,765,654 |
| | | ENCFF000QDY | Single end | 36 | hg38 | 27,090,813 | 72.89% | |
| | POLR2A phosphoS2 | ENCFF000YXC | Single end | 36 | hg38 | 24,693,516 | 84.29% | 46,902,300 |
| | | ENCFF000YXE | Single end | 36 | hg38 | 22,208,784 | 84.98% | |
| | POLR2A phosphoS5 | ENCFF000QDG | Single end | 36 | hg38 | 9,979,739 | 74.11% | 33,609,510 |
| | | ENCFF000QDO | Single end | 36 | hg38 | 23,629,771 | 72.25% | |
| Chromatin state | FAIRE-seq | GSM864361 | Single end | 35 | hg38 | 38,953,161 | 80.08% | 80,332,593 |
| | | | Single end | 35 | hg38 | 41,379,432 | 77.51% | |
| | DNase-seq (DUKE) | ENCFF000SWU | Single end | 36 | hg38 | 75,560,415 | 73.64% | 339,776,744 |
| | | ENCFF000SXA | Single end | 50 | hg38 | 128,276,304 | 70.43% | |
| | | ENCFF000SWY | Single end | 50 | hg38 | 135,940,025 | 71.27% | |
| | DNase-seq (UW) | ENCFF448LJP | Single end | 36 | hg38 | 180,112,933 | 67.09% | 199,897,086 |
| | | ENCFF810RGM | Single end | 36 | hg38 | 19,784,153 | 69.07% | |
| | ATAC-seq | GSM1782764 | Single end | 50 | hg38 | 43,114,291 | 75.82% | 73,002,093 |
| | | GSM1782765 | Single end | 50 | hg38 | 29,887,802 | 83.55% | |

**Supplementary Table 15.** Histone mark, RNA polymerase II and chromatin state datasets in GM12878

| Feature | Type | Accession in ENCODE/GEO datasets | Layout | Read length | Assembly | Uniquely mapped reads number | Uniquely mapped reads percentage | Total read number |
|---|---|---|---|---|---|---|---|---|
| Active histone marks | H3K4me1 | GSM1233902 | Paired end | 101 X 2 | hg38 | 47,598,788 | 93.95% | 127,045,430 |
| | | GSM1233903 | Paired end | 101 X 2 | hg38 | 37,904,052 | 93.08% | |
| | | GSM1233903 | Paired end | 101 X 2 | hg38 | 41,542,590 | 93.91% | |
| | H3K4me3 | ENCFF825QGB | Single end | 76 | hg38 | 31,210,351 | 89.77% | 73,139,403 |
| | | ENCFF598WCX | Single end | 76 | hg38 | 41,929,052 | 88.79% | |
| | H3K27ac | ENCFF000ASP | Single end | 51 | hg38 | 7,146,357 | 73.01% | 17,330,414 |
| | | ENCFF000ASU | Single end | 51 | hg38 | 10,184,057 | 37.71% | |
| Repressive histone marks | H3K9me3 | ENCFF000AUK | Single end | 36 | hg38 | 13,980,361 | 67.72% | 36,365,601 |
| | | ENCFF000AUO | Single end | 36 | hg38 | 11,511,369 | 63.48% | |
| | | ENCFF000AUP | Single end | 36 | hg38 | 10,873,871 | 58.63% | |
| | H3K27me3 | ENCFF000ASV | Single end | 51 | hg38 | 7,752,120 | 75.25% | 21,721,045 |
| | | ENCFF000ASZ | Single end | 51 | hg38 | 13,968,925 | 48.58% | |
| RNA polymerase II | POLR2A | ENCFF000RPI | Single end | 33 | hg38 | 16,434,825 | 70.90% | 31,063,152 |
| | | ENCFF000RPL | Single end | 33 | hg38 | 14,628,327 | 82.13% | |
| | POLR2A phosphoS2 | ENCFF000WCR | Single end | 36 | hg38 | 18,369,473 | 71.90% | 34,028,448 |
| | | ENCFF000WCS | Single end | 36 | hg38 | 15,658,975 | 78.29% | |
| | POLR2A phosphoS5 | ENCFF000OAS | Single end | 36 | hg38 | 27,190,120 | 60.19% | 56,433,145 |
| | | ENCFF000OAU | Single end | 36 | hg38 | 29,243,025 | 66.83% | |
| Chromatin state | FAIRE-seq | ENCFF000TIJ | Single end | 33 | hg38 | 45,592,393 | 76.44% | 45,592,393 |
| | DNase-seq (DUKE) | ENCFF000SLF | Single end | 20 | hg38 | 11,401,013 | 68.34% | 284,525,335 |
| | | ENCFF000SLG | Single end | 20 | hg38 | 47,604,105 | 68.41% | |
| | | ENCFF000SLL | Single end | 20 | hg38 | 104,026,587 | 71.99% | |
| | | ENCFF000SLP | Single end | 20 | hg38 | 59,238,468 | 71.91% | |
| | | ENCFF000SLR | Single end | 20 | hg38 | 62,255,162 | 71.18% | |
| | DNase-seq (UW) | ENCFF641ZDH & ENCFF306IJB | Paired end | 36 X 2 | hg38 | 186,703,242 | 80.77% | 415,475,933 |
| | | ENCFF802HKU & ENCFF885VEJ | Paired end | 36 X 2 | hg38 | 11,397,739 | 81.17% | |
| | | ENCFF813ETP & ENCFF931XDJ | Paired end | 36 X 2 | hg38 | 13,310,269 | 76.92% | |
| | | ENCFF867DMS & ENCFF045HIA | Paired end | 36 X 2 | hg38 | 204,064,683 | 77.12% | |

**Supplementary Table 16.** (see Supplementary Table 16. Transcription factor ChIP-seq datasets from ENCODE in K562 and GM12878.xlsx)

**Supplementary Table 17.** ENCODE clustered data and ChromHMM

| File Name | Assembly | Source | Link |
|---|---|---|---|
| ENCODE TFBS clusters v3 | hg19 | UCSC genome browser | http://hgdownload.soe.ucsc.edu/goldenPath/hg19/database/ |
| Broad ChromHMM | hg19 | UCSC genome browser | https://genome.ucsc.edu/ENCODE/downloads.html |
